# Deciphering of *Gorilla gorilla gorilla* Immunoglobulin Loci in Multiple Genome Assemblies and Enrichment of IMGT Resources

**DOI:** 10.1101/2024.08.19.608532

**Authors:** Chahrazed Debbagh, Géraldine Folch, Joumana Jabado-Michaloud, Véronique Giudicelli, Sofia Kossida

## Abstract

Through the analysis of immunoglobulin genes at the IGH, IGK, and IGL loci from four *Gorilla gorilla gorilla* genome assemblies, IMGT^®^ provides an in-depth overview of these loci and their individual variations in a species closely related to humans. The similarity between gorilla and human IG gene organization allowed the assignment of gorilla IG gene names based on their human counterparts. This study revealed significant findings, including variability in the IGH locus, the presence of known and new copy number variations (CNVs), and the accurate estimation of IGHG genes. The IGK locus displayed remarkable homogeneity and lacked the gene duplication seen in humans, while the IGL locus showed a previously unconfirmed CNV in the J-C cluster. The curated data from these analyses, available on the IMGT website, enhance our understanding of gorilla immunogenetics and provide valuable insights into primate evolution.

## 1 Introduction

Immunoglobulins (IG) and T-cell receptors (TR) are two types of antigen receptors, that are responsible for the extraordinary specificity and memory for antigen recognition and binding, which characterize the adaptive immune response (1–3). Immunoglobulins consist of two types of chains: heavy chains (IGH), and light chains (Kappa (IGK), or Lambda (IGL)) (4), that are encoded by four types of genes: variable (V), diversity (D), junction (J), and constant (C) (5,6).

IG genes which are distributed along the three IG loci; IGH, IGK and IGL (7) (localized on three different chromosomes in human and other vertebrates) belong to multigene families and are characterized by a high level of allelic polymorphism and a great diversity, as for example for the D genes exclusively found in the IG heavy chain (4,6,8). Moreover, IG V, D, J genes comprise specific motifs in their genomic sequences, such as recombination signals (RS), which are responsible for generating the combinatorial diversity of the variable domains. Owing to their genetic complexity (1,4), these genes are challenging to analyze and classify. Additionally, structural variations of the IG loci were shown between individuals of a same species.

IMGT^®1^, the international ImMunoGeneTics^®^ information system (9,10), established in 1989, is a high-quality integrated knowledge resource that manages sequences from genome to proteome, and structural data for immunoglobulins and T cell receptors in human and other jawed vertebrates (11,12). IMGT provides resources (database, tools, IMGT Repertoire as well as IG and TR genes and alleles reference sets) for jawed vertebrates for the analysis and understanding of immunogenetics.

Non-human primates are of great interest in comparative studies and biomedical research, due to their close similarities to human species (13). Several studies have explored the relationship between non-human primate evolution and human diseases, focusing particularly on segmental duplications found in great apes and human, which are thought to play an important role in human susceptibility to diseases, such as the case of their impact on genes associated with Mendelian diseases (14,15). Certain genes, alleles, and proteins could be implicated in causing diseases in human, yet they may be associated with normal phenotypes in gorillas. For instance, this is observed in cases such as Moyamoya disease causing deafness in human (16), as well as instances of dementia and hypertrophic cardiomyopathy (17). These studies do not include an investigation of the adaptive immune system of gorillas, nor do they incorporate genomic data related to their immunoglobulins.

Indeed, in the study of the genetics of the adaptive immune system, IMGT has been engaged for over three decades in deciphering and characterizing IG and TR loci across various species of jawed vertebrates. A recent example is the Rhesus monkey, a non-human primate, as documented in (18). We utilized several genome assemblies of *Gorilla gorilla gorilla* (Western lowland gorilla) available in the NCBI repository, which were derived from different individuals and sequencing technologies, to establish the gene repertoire of the three IG loci. Four assemblies, corresponding to three individuals, were selected based on IMGT criteria.

Our analysis of IG loci in three *Gorilla gorilla gorilla* (we will refer to ‘gorilla’ by its abbreviated name throughout this article) individuals, aiming to identify significant similarities and some differences with human, and between gorilla individuals, provides insights into human evolutionary changes in their ability to fight infections and establish an appropriate immune system (19). Additionally, it offers detailed information on the evolutionary origin of immunoglobulins and the phylogenetic relationships between primate species (17,20). The gorilla is a protected species that cannot be utilized as an animal model directly. However, knowledge of its genome is critical in immunological disease research and treatment developments.

## 2 Methods

The annotation of the IG loci was performed according to the IMGT biocuration pipeline, as previously described (21). Gorilla IG genomic sequences were analyzed and annotated by comparison with the IMGT human reference sequences ^2^.

### 2.1 Gorilla gorilla gorilla assembly selection

Seven assemblies of *Gorilla gorilla* species were available in October 2020 on NCBI (22,23), the Western lowland gorilla being the most genomic sequenced subspecies of gorillas. All of them were evaluated, and two, Kamilah_GGO_v0 (GenBank Assembly ID: GCA_008122165.1) (24) which was labelled “representative genome” of the *Gorilla gorilla gorilla* at NCBI, and Susie3 (GenBank Assembly ID: GCA_900006655.3) (25), were chosen for the quality of their IG loci which fulfilled the standard IMGT criteria for assembly selection^3^. In addition, two new *Gorilla gorilla gorilla* assemblies of the same individual, KB3781, were added in NCBI assembly (NCBI Datasets since 2024 (26)): NHGRI_mGorGor1-v1.1-0.2.freeze_mat (GenBank Assembly ID: GCA_028885495.1), the maternal haplotype, and NHGRI_mGorGor1-v1.1-0.2.freeze_pat (GenBank Assembly ID: GCA_028885475.1), the paternal haplotype (27) (Figure 1). The analysis of these four assemblies, which are all characterized by a “Chromosome” assembly level (see the Glossary^4^ of NIH), is incorporated in this study. In 2023, The NHGRI_mGorGor1-v1.1-0.2.freeze_pri assembly (GenBank Assembly ID: GCA_029281585.1), the principal haplotype of KB3781, became the “representative genome” of the *Gorilla gorilla* species (it includes the maternal autosomes, unplaced sequence identified as maternal, chrX, chrY, and MT), and the three assemblies of the KB3781 genome were updated in 2024 (Figure 1). As for the two assemblies of the Kamilah individual published in 2023 Kamilah_GGO_hifiasm-v0.15.2.pri (GenBank Assembly ID: GCA_030174185.1), and Kamilah_GGO_hifiasm-v0.15.2.alt (GenBank Assembly ID: GCA_030174155.1) (28), the corresponding biocuration results are solely presented in the discussion for reasons that will get apparent further down in the manuscript.

**Figure 1.**
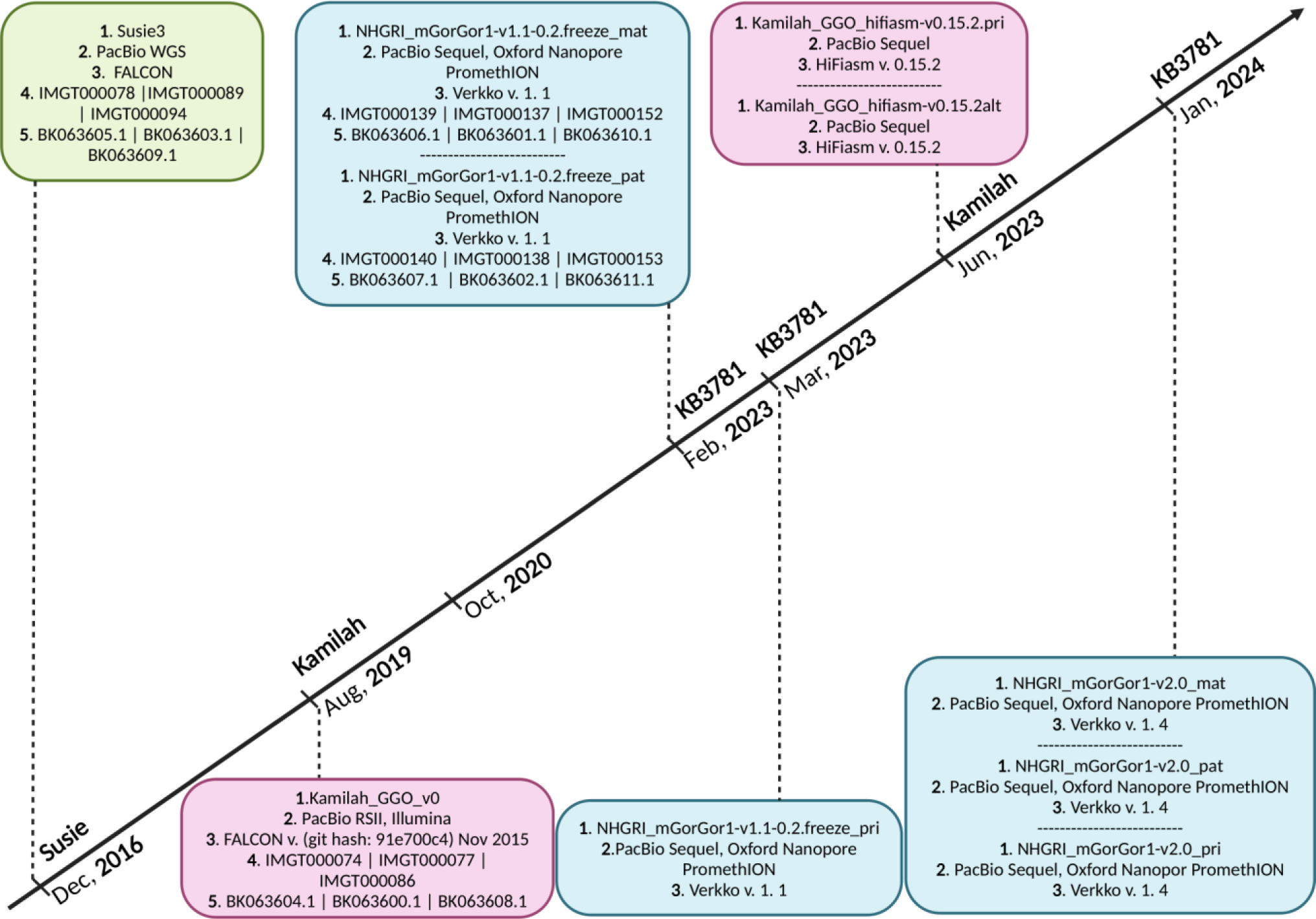
Timeline of *Gorilla gorilla gorilla* genome assemblies published on NCBI, including: green box: assembly of Susie individual, pink boxes: assemblies of Kamilah individual, blue boxes: assemblies of KB3781 individual, 1. assembly name 2. sequencing technology, 3. assembly method, 4. IMGT annotation accession number, and 5. NCBI Third Party Annotation (TPA) accession number. The *Gorilla gorilla gorilla* IMGT annotation project of IG loci started October 2020. It currently incorporates assemblies at chromosome level of Susie, Kamilah and KB3781 individuals, published in December 2016, August 2019 and February 2023, respectively. The principal assembly of KB3781 individual was published in March 2023. The two new assemblies (principal and alternate) of Kamilah, that are available at contig level, were published in June 2023. In January 2024, new versions for the three assemblies of KB3781 individuals were published.

### 2.2 Loci sequence extraction from NCBI and integration in IMGT

For each assembly, the localization of the three IG loci (IGH, IGK and IGL) on chromosomes was determined by comparison to the IMGT human IG reference set, by using BLAST (29). The delimitations of the IGK and IGL loci were defined by the identification of the flanking non IG genes which are conserved among species upstream of the first IG gene and downstream of the last IG gene, called “IMGT bornes^5^” (30). If and only if the distance of the “IMGT bornes” is over 10.000 bp from the first and the last IG gene in 5’ and in 3’ respectively, the delimitations of the IGK and IGL loci are defined by 10.000 bp upstream of the first IG gene and 10.000 bp downstream of the last IG gene. Due to the absence of “IMGT bornes” for IGH loci, the gorilla IGH loci were delimitated by 10.000 bp upstream of the first IG gene and 11.000 bp (exclusively for gorilla IGH locus) downstream of the last IG gene. The corresponding nucleotide sequences were extracted from the NCBI chromosome sequences, and IMGT/LIGM-DB (31) entries were created.

### 2.3 V, D, J and C genes annotation

The V, D, J and C genes were first detected and delimitated along the IMGT/LIGM-DB (31) genomic sequences (IGH, IGK and IGL loci), with IMGT/LIGMotif (32). IG genes were characterized and classified using alignments by BLAST (29), Clustal Omega (33), and by implementing the IMGT unique numbering (34,35), and annotation rules of the IMGT Scientific chart, based on the IMGT-ONTOLOGY concepts (36) for the genes and alleles functionalities^6^ and the setting of gorilla IG genes nomenclature^7^. Due to the extremely high sequence similarity between gorilla and human (37), which was confirmed at the level of the IG loci in the early steps of biocuration (see section ‘4 Results’ and supplementary Tables 1-3), gorilla IG genes were named according to their human counterpart based on their sequence similarity and their position in the locus (nomenclature by orthology). Additional genes in gorilla loci compared to the human species were named as inserted genes (incrementation of the number of the V gene sub-positions from 3’ to 5’, and the number of the D gene sub-positions from 5’ to 3’, and addition of the Latin alphabet letters, from 5’ to 3’ for J and C genes), another case of additional genes, the genes were named as duplicated (name of the “initial” gene, with addition of the letter “D”). IG genes were integrated into IMGT/GENE-DB (38) and the synthesis of biocuration data regarding the loci, genes, alleles and proteins into IMGT Web resources for IG^8^.

The NCBI Third Party Annotation (TPA^9^) (39) accession numbers were provided for the three *Gorilla gorilla gorilla* IG loci.

### 2.4 CNV characterization

The names of Copy Number Variation (CNV) in gorilla IG loci are identical to those of human counterparts (30), if equivalent. For gorilla potential specific CNV, they are provisionally named CNVp, plus an incremental number.

## 3 Results

The three loci IGH, IGK, and IGL of genes were extracted from four *Gorilla gorilla gorilla* genome assemblies of three individuals publicly available at NCBI, and annotated according to IMGT standards. Table 1 summarizes loci information and gives total number of genes four each assembly, and their accession numbers initially created in IMGT/LIGM-DB (31), and the ones assigned in the TPA database (39).

**Table 1.**
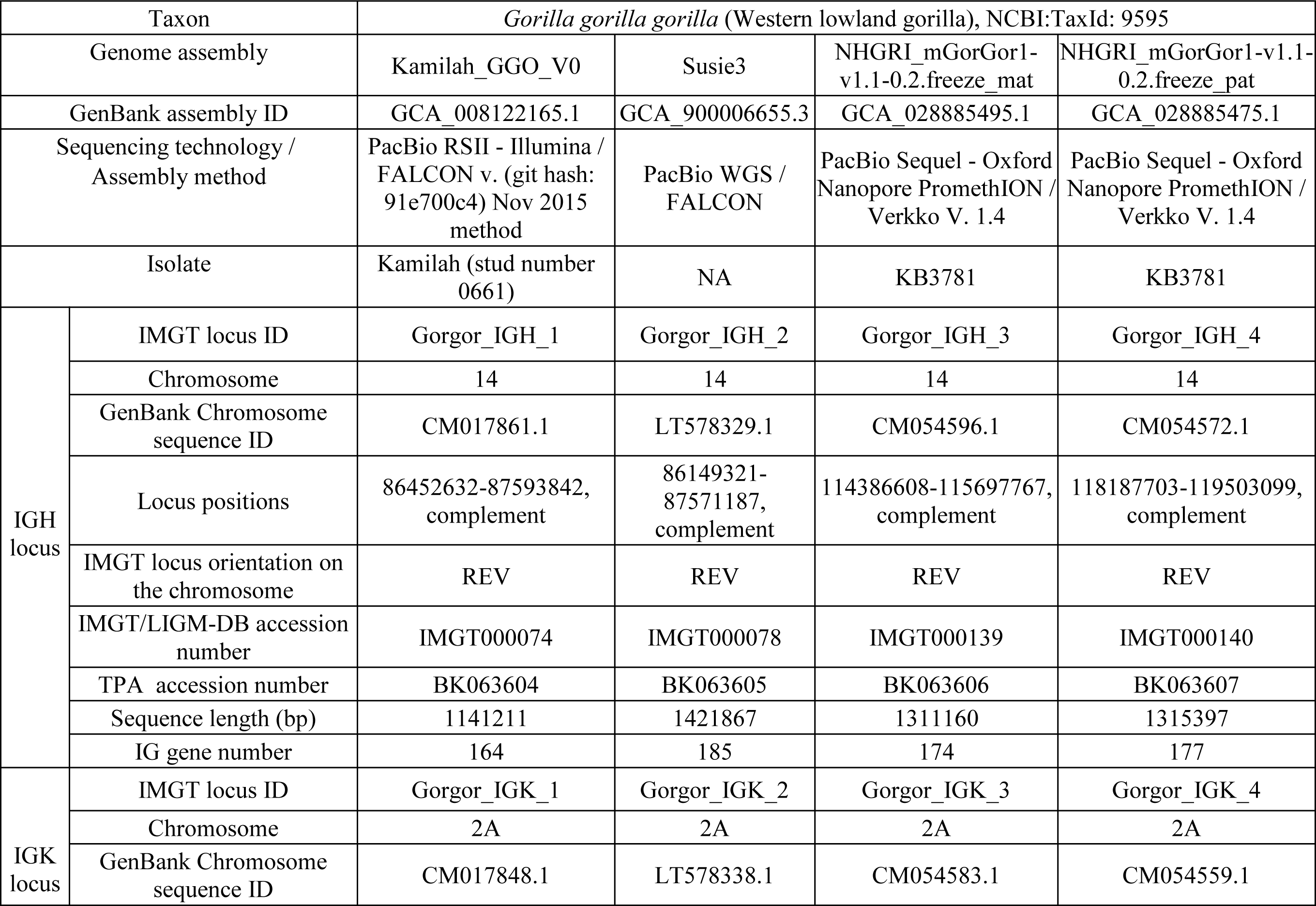
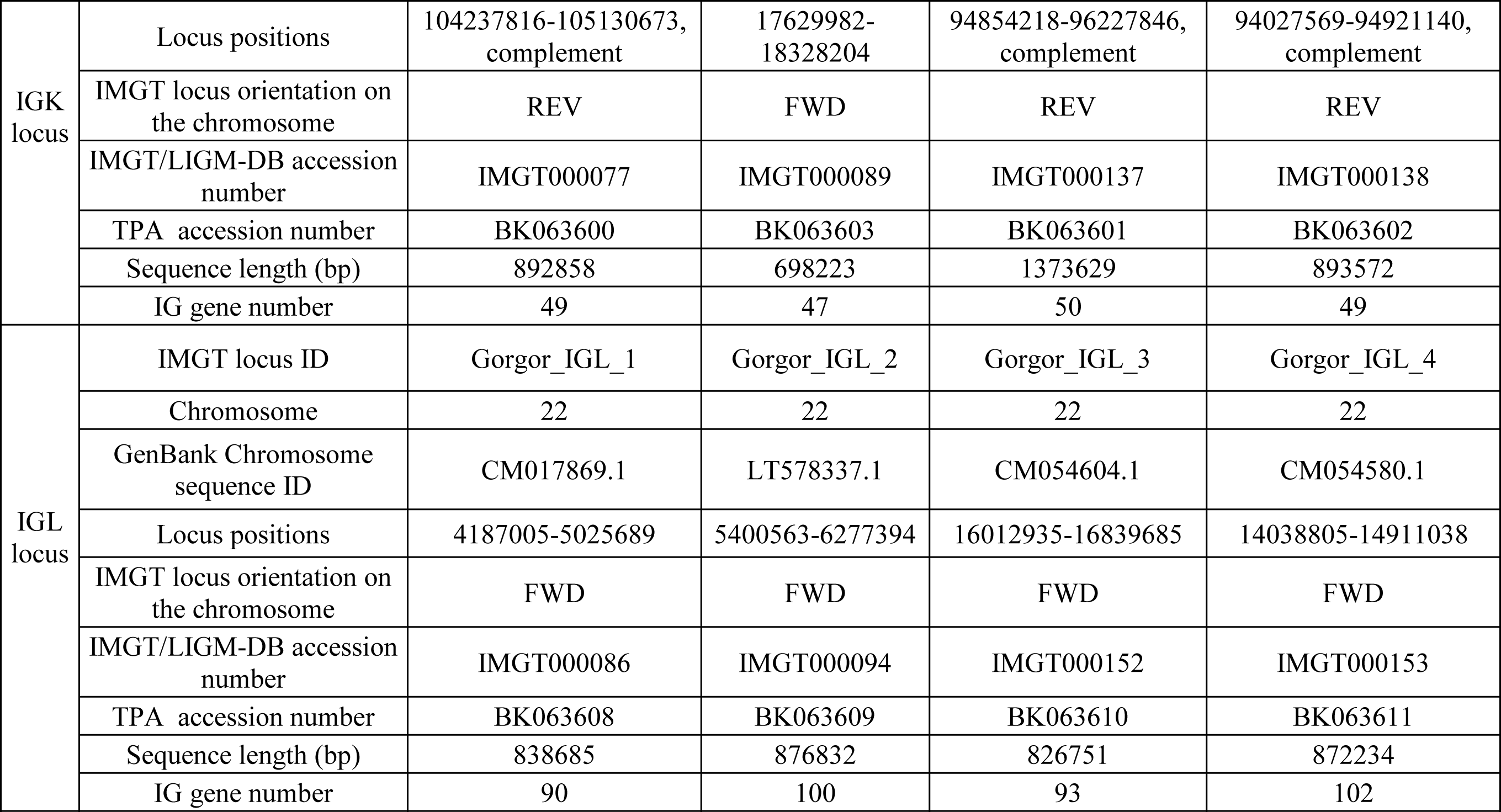
Information about genome assembly and IGH, IGK and IGL loci for the four *Gorilla gorilla gorilla* assemblies.

| Taxon |  | *Gorilla gorilla gorilla* (Western lowland gorilla), NCBI:TaxId: 9595 |  |  |  |
| --- | --- | --- | --- | --- | --- |
| Genome assembly |  | Kamilah_GGO_V0 | Susie3 | NHGRI_mGorGor1-v1.1-0.2.freeze_mat | NHGRI_mGorGor1-v1.1-0.2.freeze_pat |
| GenBank assembly ID |  | GCA_008122165.1 | GCA_900006655.3 | GCA_028885495.1 | GCA_028885475.1 |
| Sequencing technology /<br>Assembly method |  | PacBio RSII - Illumina /<br>FALCON v. (git hash:<br>91e700c4) Nov 2015<br>method | PacBio WGS /<br>FALCON | PacBio Sequel - Oxford<br>Nanopore PromethION /<br>Verkko V. 1.4 | PacBio Sequel - Oxford<br>Nanopore PromethION /<br>Verkko V. 1.4 |
| Isolate |  | Kamilah (stud number<br>0661) | NA | KB3781 | KB3781 |
| IGH<br>locus | IMGT locus ID | Gorgor_IGH_1 | Gorgor_IGH_2 | Gorgor_IGH_3 | Gorgor_IGH_4 |
|  | Chromosome | 14 | 14 | 14 | 14 |
|  | GenBank Chromosome<br>sequence ID | CM017861.1 | LT578329.1 | CM054596.1 | CM054572.1 |
|  | Locus positions | 86452632-87593842,<br>complement | 86149321-87571187,<br>complement | 114386608-115697767,<br>complement | 118187703-119503099,<br>complement |
|  | IMGT locus orientation on<br>the chromosome | REV | REV | REV | REV |
|  | IMGT/LIGM-DB accession<br>number | IMGT000074 | IMGT000078 | IMGT000139 | IMGT000140 |
|  | TPA accession number | BK063604 | BK063605 | BK063606 | BK063607 |
|  | Sequence length (bp) | 1141211 | 1421867 | 1311160 | 1315397 |
|  | IG gene number | 164 | 185 | 174 | 177 |
| IGK<br>locus | IMGT locus ID | Gorgor_IGK_1 | Gorgor_IGK_2 | Gorgor_IGK_3 | Gorgor_IGK_4 |
|  | Chromosome | 2A | 2A | 2A | 2A |
|  | GenBank Chromosome<br>sequence ID | CM017848.1 | LT578338.1 | CM054583.1 | CM054559.1 |
| IGK locus | Locus positions | 104237816-105130673, complement | 17629982-18328204 | 94854218-96227846, complement | 94027569-94921140, complement |
|  | IMGT locus orientation on the chromosome | REV | FWD | REV | REV |
|  | IMGT/LIGM-DB accession number | IMGT000077 | IMGT000089 | IMGT000137 | IMGT000138 |
|  | TPA accession number | BK063600 | BK063603 | BK063601 | BK063602 |
|  | Sequence length (bp) | 892858 | 698223 | 1373629 | 893572 |
|  | IG gene number | 49 | 47 | 50 | 49 |
| IGL locus | IMGT locus ID | Gorgor_IGL_1 | Gorgor_IGL_2 | Gorgor_IGL_3 | Gorgor_IGL_4 |
|  | Chromosome | 22 | 22 | 22 | 22 |
|  | GenBank Chromosome sequence ID | CM017869.1 | LT578337.1 | CM054604.1 | CM054580.1 |
|  | Locus positions | 4187005-5025689 | 5400563-6277394 | 16012935-16839685 | 14038805-14911038 |
|  | IMGT locus orientation on the chromosome | FWD | FWD | FWD | FWD |
|  | IMGT/LIGM-DB accession number | IMGT000086 | IMGT000094 | IMGT000152 | IMGT000153 |
|  | TPA accession number | BK063608 | BK063609 | BK063610 | BK063611 |
|  | Sequence length (bp) | 838685 | 876832 | 826751 | 872234 |
|  | IG gene number | 90 | 100 | 93 | 102 |

The resulting data for locus, assemblies, gene, allele, sequence, protein, expression cDNA, and statistics is available in IMGT Web resources, and the detailed list is provided in Supplementary Table 4.

The gorilla IG gene names were assigned by orthology with human and according to IMGT gene nomenclature principles. The percentages of identity between the closest gorilla alleles and their human counterpart are reported in Supplementary Tables 1-3. The loci and genes data from Kamilah_GGO_v0 were chosen as reference for the analysis of loci variations between gorilla individuals in terms of gene and allele content. The IMGT gene order of V, D, J, C and non-IG genes from 5’ to 3’ for each locus, was initially established for the Kamilah_GGO_v0 assembly. According to this, the gene order of additional genes in the three other assemblies was identified (Supplementary Tables 1-3).

Figure 2 presents an overview of the number of annotated IG genes that are common or unique within the four gorilla assemblies. The locus IGH appears to be more variable in terms of common genes (73% of total annotated IGH genes) across the four assemblies, compared to the IGK (94% of total common annotated IGK genes) and IGL (84% of total common annotated IGL genes) loci, which are more conserved among the three individuals. The heterogeneity of the IGH locus between the four gorilla assemblies is particularly evident in the detected CNVs, as well as the duplication and insertion of genes throughout the locus (see section ‘3.1 *Gorilla gorilla gorilla* IGH locus’).

**Figure 2.**
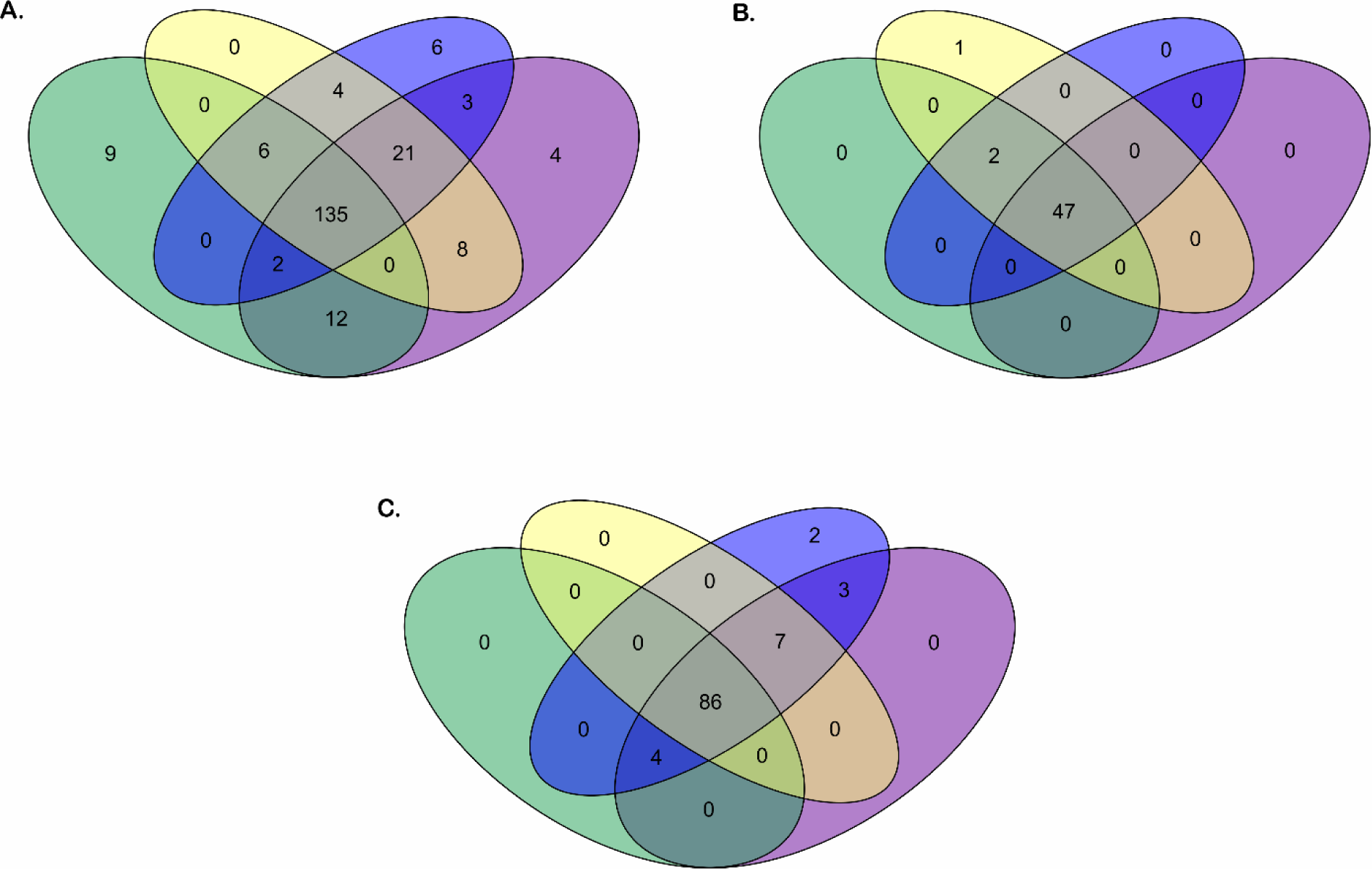
Green oval: Kamilah_GGO_v0 assembly. Purple oval: Susie3 assembly. Yellow oval: NHGRI_mGorGor1-v1.1-0.2.freeze_mat assembly. Bleu oval: NHGRI_mGorGor1-v1.1-0.2.freeze_pat assembly. The Venn diagrams (A), (B) and (C) show respectively the common or unique IGH, IGK and IGL genes between the four *Gorilla gorilla gorilla* assemblies.

### 3.1 Gorilla gorilla gorilla IGH locus

#### 3.1.1 Localization and description of IGH locus

The IGH locus of gorilla extends from 10 kb upstream of the most 5’ gene in the locus, IGHV(III)-82, to 11 kb downstream of the most 3’ gene, IGHA2. It comprises between 164 and 185 genes depending on the assembly and all of them are in the sense orientation in the locus (Table 1). According to the description and annotation of the locus with “IMGT Labels^10^” (40), the IGH locus is composed of four clusters of the same gene type: 120 to 135 V genes (V-CLUSTER), 18 to 32 D genes (D-CLUSTER), 8 to 9 J genes (J-CLUSTER) and 4 to 13 C genes (C-CLUSTER). Its organization is very close to the human one. Interestingly, the eight known RPI (Related proteins of the immune system) genes within the human IGH locus were also identified in gorilla assemblies (Table 1, Supplementary Figures 1-4, Supplementary Table 1).

#### 3.1.2 IG gene organization in the gorilla IGH locus

##### 3.1.2.1 IGHV genes cluster

Overall, 157 IGHV genes and 316 IGHV alleles were identified in the gorilla IGH loci of the four assemblies (Figure 2). Based on their high level of sequence similarity with human IGH genes, 105 gorilla V genes could be classified into 8 subgroups and 52 others in three clans.

A phylogenetic tree was built from a sequence set including the first allele (the reference sequence of each gorilla gene) and the first allele for human genes. This phylogenetic tree was created to highlight the close similarity of genes between both species within a subgroup (Figure 3). The pseudogenes of the clan IGHV(III) intercalates with the subgroup IGHV3, because of the sequence similarity and according to the “IMGT IGH clan tree^11^”. It shows that the gorilla subgroup/clan genes are grouped in the same branch with the corresponding human gene.

**Figure 3.**
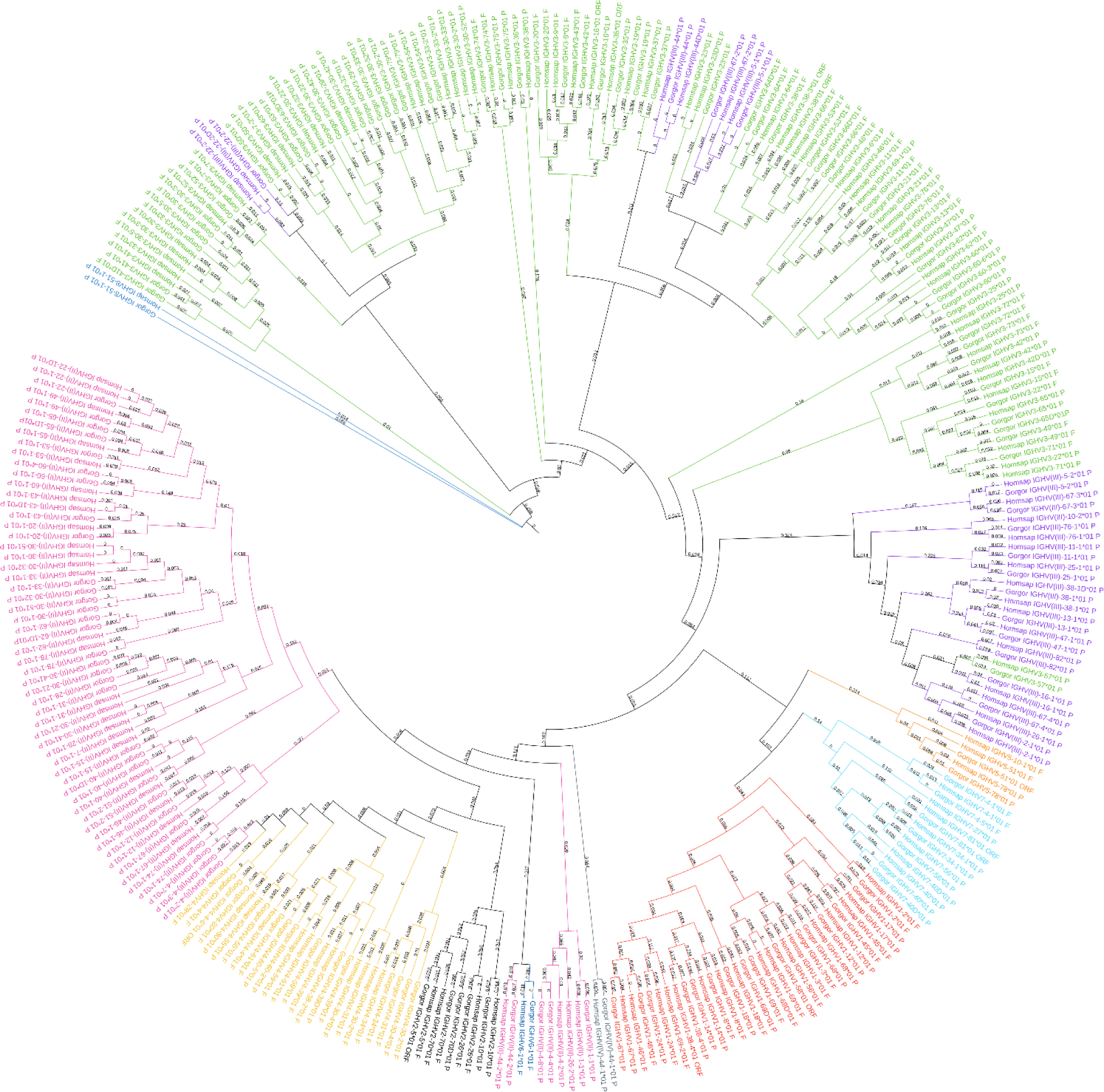
Phylogenetic tree of all IGH subgroups and clans for *Gorilla gorilla gorilla* and *Homo sapiens*, using first allele of each gene. The different colors highlight the different subgroups and clans. Tree generated using NGPhylogeny.fr (41) and iTOL v6 (42).

A total of 136 gorilla IGHV genes were named according to their human counterpart (Supplementary Table 1. The names of 29 additional IGHV genes, only present in gorilla genome (underlined in green in the supplementary Table 1) were set by applying IMGT nomenclature for inserted genes, or by taking into account the evidence gene bloc duplication: the two duplicated blocs including the genes IGHV3-41D to IGHV4-39D, and the genes IGHV(II)-62-1D to IGHV3-66D, which show over 99,5% of identity with the initial blocs IGHV3-41 to IGHV4-39 and IGHV(II)-62-1 to IGHV3-66, respectively.

Based on this comparative approach, we also identified 29 human IGHV genes for which the gorilla counterpart cannot be found in the IGH locus in any of the four assemblies (underlined in yellow in the Supplementary Table 1). Interestingly, all of them (except IGHV7-77) are located within well-known human CNVs (30).

Figure 4 shows number of genes within subgroups and clans. Interestingly the majority of functional genes belong to IGHV3 subgroup (Supplementary Table 5), as in human. The human IGHV3 genes are known to be selected in response to superantigens (43,44). The expansion of this subgroup IGHV3, could also be results of the major CNV3 and the gorilla potential CNVp1. The detailed list of alleles per IGHV subgroup or clan and per functionality present in each assembly can be obtained from the section “Locus gene repertoire per IMGT annotated assembly^12^” of IMGT Web resources (after selection of the species and the locus).

**Figure 4.**
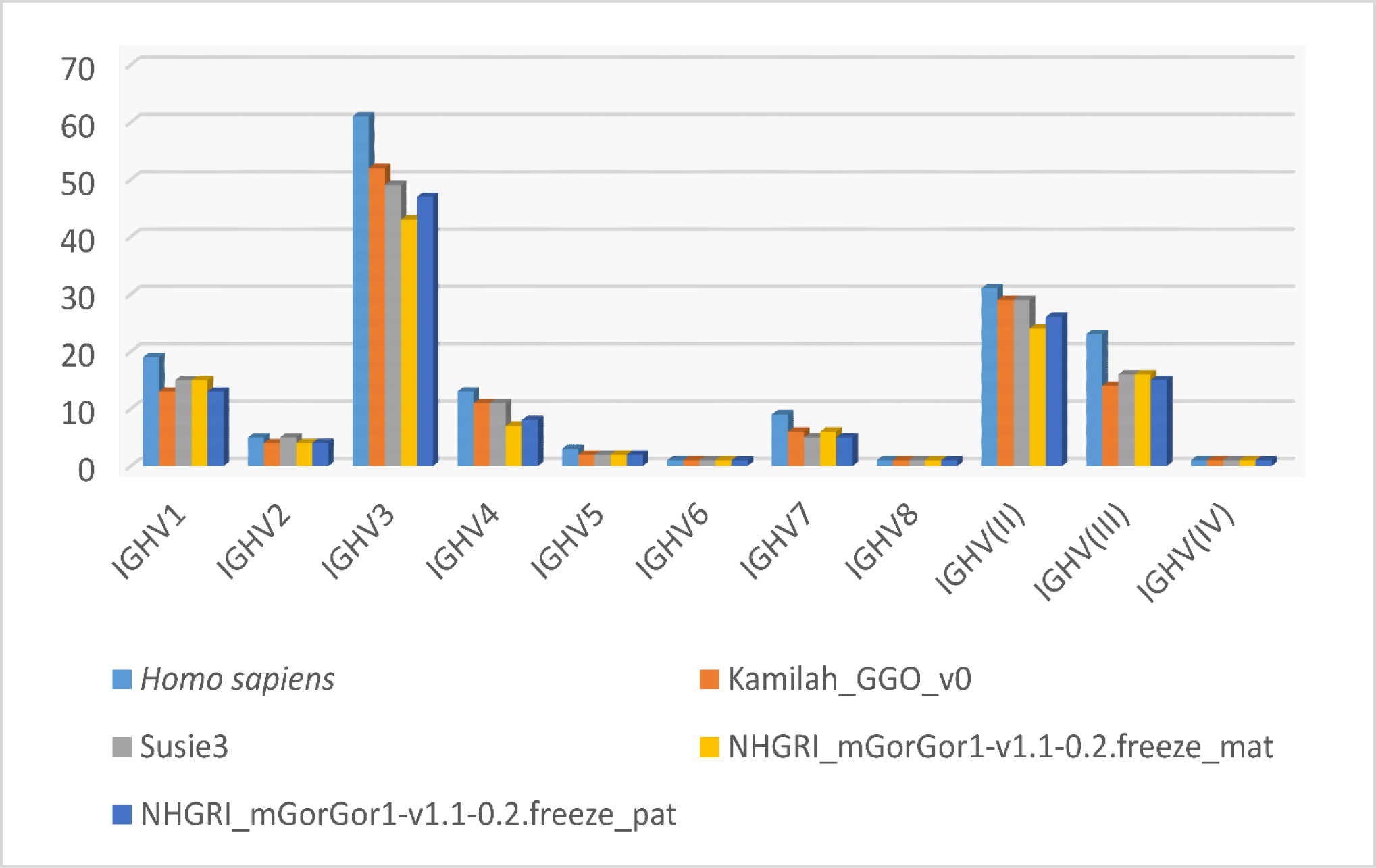
Number of IGHV genes per IMGT subgroup/clan in human and in the four assemblies of *Gorilla gorilla gorilla*.

##### 3.1.2.2 IGHD and IGHJ genes clusters

The analysis of the gorilla IGH locus of four assemblies allowed the identification of 32 IGHD genes classified in seven sets as for human and 43 IGHD alleles of which 29 are functional (Supplementary Table 1). Fifteen consecutive genes (IGHD6-5-1, IGHD1-5-2 IGHD2-5-3, IGHD3-5-4, IGHD4-5-5, IGHD5-5-6, IGHD6-6, IGHD1-7, IGHD2-8, IGHD3-10, IGHD4-11, IGHD5-12, IGHD6-13 and IGHD1-14) are missing in Kamilah_GGO_V0 assembly due to a 20 kb gap within the D-CLUSTER area (Supplementary Figure 1). However, all IGHD genes were recovered in the new assemblies of Kamilah’s genome (data not shown, see section ‘4.5 Assemblies of “Kamilah” individual’). Among them, 26 have been named according to their human counterpart. Additionally, six new IGHD genes were identified (GHD6-5-1, IGHD1-5-2 IGHD2-5-3, IGHD3-5-4, IGHD4-5-5, IGHD5-5-6), they might be gorilla specific, since human counterparts were not found in human IGH locus whatever the 16 assemblies processed by IMGT. Based on the results obtained from the four assemblies, it appears that the gorilla IGH locus does not include an ortholog for the human IGHD3-9.

Nine IGHJ genes and 16 alleles have been identified and localized in gorilla IGH locus. The IGHJ genes organization is comparable to the one of the human J-CLUSTER. They were classified into six subsets and all of them were named according to their human counterparts. The gorilla IGHJ2 gene was not found in the Kamilah_GGO_v0 assembly. It is also missing in the recent assembly version of this genome, Kamilah_GGO_hifiasm-v0.15.2.pri, but it is present in Kamilah_GGO_hifiasm-v0.15.2.alt (data not shown, see section ‘4.5 Assemblies of “Kamilah” individual’). Therefore, this variation could correspond to a CNV in the gorilla IGH locus.

##### 3.1.2.3 IGHC genes cluster

A total of 13 gorilla IGHC genes and 29 alleles were identified taken into account the four assemblies (Supplementary Table 1; Supplementary Figures 1-4; Dynamic gene tables per IMGT group and per species^13^). These genes code for the five isotype classes: IgM (IGHM), IgD (IGHD), IgG (IGHG1, IGHG2, IGHG3A, IGHG3B, IGHG3C, IGHG4, IGHGP), IgE (IGHEP1, IGHE), and IgA (IGHA1, IGHA2).

Interestingly, the IGHG genes, to our knowledge, are being comprehensively characterized for the first time. Previous studies relied on a single assembly, Kamilah_GGO_v0, which included only four IGHC genes: IGHM, IGHD, IGHG3A, and IGHG1 (45,46).

The gorilla IGH locus includes three IGHG3 genes which differ from each other by their number of hinge exons (two or five for IGHG3A depending on the allelic polymorphism, two for IGHG3B, and four for IGHG3C). The gorilla C-CLUSTER shows a highly similar organization to the human one. The main difference is the addition of two IGHG3 genes, presumably IGHG3A and IGHG3B since the human IGHG3 and the gorilla IGHG3C share the same number of hinge exons and almost 100% of identity in all their exon sequences.

### 3.2 Gorilla gorilla gorilla IGK locus

#### 3.2.1 Localization and description of IGK locus

The gorilla IGK locus has a reverse orientation (REV) on the chromosome 2A for the three assemblies Kamilah_GGO_v0, NHGRI_mGorGor1-v1.1-0.2.freeze_mat and NHGRI_mGorGor1-v1.1-0.2.freeze_pat, whereas the Susie3 IGK locus is forward (FWD) (Table 1). However, the REV or FWD orientation does not appear to affect the genomic structure, gene organization, or gene functionality. According to IMGT rules for quality assessment of IG and TR loci in genome assemblies, the IGK locus was satisfactory for genomic annotation across all four assemblies.

Two IMGT flanking genes “IMGT bornes” (30) were identified for the IGK locus delimitation of many species (see the section “Locus bornes: IGK locus 5’ and 3’ bornes^14^” of IMGT Repertoire), including the gorilla. The 5’ end IMGT flanking gene, “IMGT borne”, PAX8 (NCBI Gene ID: 101137174) is 125 kb upstream of IGKV1-49, the most 5’ gene in the locus, and the 3’ end IMGT flanking gene, “IMGT borne”, RPIA (NCBI Gene ID: 101148377) is 200 kb downstream of IGKC, the most 3’ gene in the locus. “IMGT bornes” were identified in the four annotated assemblies.

The IGK locus extends from 10 kb upstream, IGKV1-49, to 10 kb downstream, IGKC. The differences of IGK locus size observed for the four assemblies (varying from 698 kb to 1374 kb) set in the area between the IGKV2-29 and IGKV3-31: this part of the locus varies in length and seems not comprise IGK genes or any other genes. The IGK locus comprises a cluster of 41 to 44 V genes (V-CLUSTER) for a large part and a cluster of 5 J and 1 C genes (J-C-CLUSTER) (Table 1, Supplementary Figures 5-8, Supplementary Table 2).

#### 3.2.2 IG gene organization in the gorilla IGK locus

##### 3.2.2.1 IGKV genes cluster

A total of 44 IGKV genes were identified from the annotation of the four assemblies. 31 of them show polymorphic alleles and in total 92 IGKV alleles were characterized (Figure 2, Supplementary Table 2).

The IGKV genes were classified into seven subgroups defined according to the IMGT-ONTOLOGY (36) and their sequence similarity with the IGKV human subgroups. The phylogenetic tree in Supplementary Figure 13, constructed from the first allele of all gorilla and human IGKV genes, displays the distances between the IGKV genes of both species. It shows that the gorilla genes are grouped on the same clade as their human counterpart genes.

The number of IGKV genes in gorilla is slightly over half that of the human IGKV gene number, with 76 localized IGKV genes in the main human locus. The human IGK locus comprises a proximal V-CLUSTER (p) of 40 IGKV genes, and a distal V-CLUSTER (d) of 36 genes, from 3’ to 5’ (1,47). The first eight gorilla IGKV genes starting from the 3’ to 5’ of the IGK locus, have an extremely close organization, gene order, sequence similarity and gene orientation in the locus (IGKV4-1 IGKV5-2 have opposite orientation in the locus, as well as in human) to that of the human proximal (p) IGK V-CLUSTER. The other gorilla IGKV genes are mostly closer to the human distal (d) IGK V-CLUSTER (Supplementary Table 2). Moreover, counterparts of the human IGKV6D-41, IGKV1D-42 and IGKV1D-43 were identified in gorilla assemblies whereas they were not in the human proximal (p) IGK V-CLUSTER. Conversely, it should also be noted that there is no counterpart of IGKV1-9 (located in the human proximal IGK V-CLUSTER) in the gorilla assemblies, nor is there a corresponding duplicated gene in the human distal IGK V-CLUSTER.

Therefore, the IGK genes nomenclature of the human proximal IGKV V-CLUSTER was assigned to all gorilla IGKV genes, and IGKV6-41, IGKV1-42 and IGKV1-43 genes, according to their orthology, and especially since gorilla does not have duplicated IGK V-CLUSTER. For additional genes, names were assigned according to IMGT nomenclature rules.

The genes between IGKV1-44 and IGKV1-49 are additional in gorilla IGK locus compared to the human one, the positional nomenclature was adopted by incrementation of the position number on the locus.

Almost 1/3 of gorilla IGKV genes (16-17) belong to IGKV1 subgroup which includes the highest number of functional genes, and the other 1/3 (13-15) belongs to IGKV2 subgroup which includes the highest number of pseudogenes (Supplementary Table 6, Supplementary Figure 15).

##### 3.2.2.2 IGKJ and IGKC genes clusters

The annotation of the four gorilla IGK assemblies allowed to highlight five IGKJ genes belonging to five sets 1, 2, 3, 4 and 5, one gene for each set, and one unique constant IGKC gene (Supplementary Table 2). In examining the assemblies, these genes appear to be minimally or not at all polymorphic: only one additional allele was shown for the IGKJ gene and none for IGKC (Figure 2).

### 3.3 Gorilla gorilla gorilla IGL locus

#### 3.3.1 Localization and description of IGL locus

The IGL locus for the four selected gorilla assemblies is delimitated by flanking genes, “IMGT bornes’’ (30) identified in other species (see the section “Locus bornes: IGL locus 5’ and 3’ bornes^15^” of IMGT Repertoire) (Table 1). The 5’ end IMGT IGL Locus borne is the TOP3B gene (NCBI Gene ID: 101141903) which is in reverse orientation, and the 3’ end IMGT IGL Locus borne gene, the RSPH14 gene (NCBI Gene ID: 101130781). The IGL locus extends from 10 kb upstream the most 5’ gene in the locus, IGLV(I)-70-1, to 10 kb downstream of the most 3’ gene in the locus, IGLC7, and comprises a cluster of 76 to 86 V genes (V-CLUSTER) and a cluster of 7 to 8 IGLJ and C genes (J-C-CLUSTER). Interestingly, six known RPI (Related proteins of the immune system) genes within the human IGL locus were also identified in gorilla assemblies (Table 1, Supplementary Figures 9-12, Supplementary Table 3).

#### 3.3.2 IG gene organization in the gorilla IGL locus

##### 3.3.2.1 IGLV genes cluster

The analysis of the four assemblies allowed the identification of 86 IGLV genes, in total. From genes annotation, 53 genes show allelic polymorphism and in total 163 IGLV alleles were characterized (Figure 2, Supplementary Table 3).

The IGLV genes were classified into eleven subgroups and seven clans defined according to IMGT-ONTOLOGY (36) and to their sequence similarity with the human IGLV subgroups. The phylogenetic tree of the Supplementary Figure 14, built from the first allele of all gorilla and human IGLV genes, displays the distance between the IGLV genes of both species and shows that the gorilla genes are grouped in the same clade with their human counterpart gene. The clans IGLV(I), IGLV(II) and IGLV(V) are interspersed between subgroups because of the sequence similarity and the gene nomenclature according to the “IMGT IGL clan tree^16^”.

As for human and other non-human primates, the gorilla IGLV3 subgroup gathers the highest number of genes (20 to 24 depending on the assembly), with approximately the same number of functional genes and pseudogenes (Supplementary Table 7, Supplementary Figure 16).

The comparison of the IGL locus from the four assemblies with the IGL locus organization of human highlights common features between the four assemblies: two blocs of human IGLV genes are absent in the gorilla locus (highlighted in yellow in Supplementary Table 3): a bloc of 7 genes (IGLV(VII)-41-1, IGLV1-41, IGLV1-40, IGLV5-39, IGLV(I)-38, IGLV5-37 and IGLV1-36) and a bloc of 5 genes (IGLV(IV)-66-1, IGLV(V)-66, IGLV(IV)-65, IGLV(IV)-64 and IGLV(I)-63). The availability of assemblies from more individuals should help to confirm if this observation corresponds to a gorilla specific feature. Another gorilla specific attribute would correspond to the 16 IGLV genes (highlighted in green in the Supplementary Table 3) present in the four gorilla assemblies but not in human. We also identify a potential CNV between IGLV3-24-2 and IGLV3-27, an area of eight IGLV genes not identified in all four assemblies.

We noticed that six genes: IGLV(I)-34-1, IGLV2-34, IGLV2-33, IGLV3-32, IGLV3-31, IGLV3-30 were not identified in the Kamilah_GGO_v0 assembly. This seems to be linked to the presence of a gap of 47 kb in this position and this cannot be considered as potential CNV.

##### 3.3.2.2 IGLJ-C genes cluster

The gorilla IGL J-C-CLUSTER is composed of seven tandems of IGLJ and IGLC genes, or eight (IGLJ2A and IGLC2A for NHGRI_mGorGor1-v1.1-0.2.freeze_pat only, which is considered as potential CNVp2 in gorilla). The IGLJ genes show a very low allelic polymorphism (only two alleles for IGLJ5).

## 4 Discussion

The identification of the gorilla IG genes and alleles, along with the characterization of their genomic organization detailed in the present study, increases our knowledge of the genetics of the adaptive immune response in jawed vertebrates. Additionally, it provides interesting clues regarding the molecular evolution and conservation of gorilla IG loci among primates, as well as the individual variations within the population.

To detect potential evolutionary events in germline DNA sequences of gorilla IG loci, we relied on gorilla-human comparative genomics study. Whatever the locus, the three individuals and the four NCBI assemblies (Table 1), the gorilla IG loci have retained a structure close to the related locus in human with approximatively the same number of genes (except for IGK if we count the number of genes in the proximal and distal copies). Sequences of both species present high similarity which is closely correlated with the taxonomic relationship. The speciation event led to the conservation of orthologous genes in gorilla, and according to IMGT gene nomenclature, IG gorilla genes were assigned the names of their orthologous human IG genes, if any. In addition, orthologous genes positions in the loci, and the use of the IMGT positional nomenclature for gorilla genes were also confirmed with the detection of flanking genes, “IMGT bornes” (30), when they exist (IGL and IGK loci), and with highly conserved Related Proteins of the Immune system (RPI) in the IGH and IGL loci. These RPI sequences are conserved in all mammals and used as markers in the locus (30).

Comparison of the gorilla IG loci from the four assemblies (Kamilah_GGO_v0, Susie3, NHGRI_mGorGor1-v1.1-0.2.freeze_mat and NHGRI_mGorGor1-v1.1-0.2.freeze_pat) highlights genomic variations that have been observed exclusively in gorilla species, which might suggest that the genome is accumulating unique variations depending on each individual. The IG genomic sequences of the four assemblies were selected according to IMGT rules for assessment of IG and TR loci in genome assemblies. It is worth mentioning that the NHGRI_mGorGor1-v1.1-0.2.freeze_pri, the NCBI “representative genome” of Western lowland gorilla since March 2023, includes exactly all IG genes and alleles of NHGRI_mGorGor1-v1.1-0.2.freeze mat (from which the IG loci were analyzed), and therefore the IG loci of this assembly are not detailed in the current article.

After analyzing all previous assemblies, two additional ones were published online; Kamilah_GGO_hifiasm-v0.15.2.pri and Kamilah_GGO_hifiasm-v0.15.2.alt, which were acquired from the same individual, Kamilah, utilizing PacBio Sequel technology and HiFiasm v. 0.15.2 assembly method. The latter two assemblies are at the contig level, therefore they were not fully analyzed and included in this article, because they are not on the chromosome level.

### 4.1 V-GENE multigene families, allelic polymorphism, gene insertion/deletion and CNVs identification in Western lowland gorilla IG loci

The diversity of the IG variable domains is partly generated by the repertoire of large numbers of variable (V) genes in the germline DNA (8). This is especially true for the V genes of the heavy chain, which are more numerous than those of light chains in most species, which is the case in Western lowland gorilla. As mentioned in (8) paper, the reason behind the expansion and contraction of the IGHV multigene families in jawed vertebrates is still poorly understood. The duplication and divergence events in IG loci are governed by different patterns driven by natural selection.

Comparison of V genes by multiple alignment between the annotated loci of the different assemblies, revealed the duplication of certain genes, as well as the divergence of duplicated genes that occur during locus evolution. In both cases, the genes are considered as phylogenetically related. The more duplicated genes that occur, the more nonfunctional genes are produced in the IGHV multigene families (6). Importantly, these pseudogenes are carefully considered in the IMGT annotation of germline DNA, as they provide precious clues of the organism’s evolution.

#### 4.1.1 IMGT Subgroups of IGH, IGK and IGL variable genes

IMGT Subgroups names of non-human primates have been assigned by homology with those of human. The classification of gorilla variable genes into IMGT subgroups highlights the abundance of subgroups: IGHV3 for IGH, IGKV1 for IGK, and IGLV3 for IGL (Figure 4, Supplementary Table 1-3, Supplementary Figure 15-16). Interestingly, this result is also observed for the other non-human primate species, such as Rhesus monkey (*Macaca mulatta*) (18), Sumatran orangutan (*Pongo abelii*), Bornean orangutan (*Pongo pygmaeus*), and even for IGH and IGL of Ring-tailed lemur (*Lemur catta*), a more distant primate species in the taxonomy classification (data available at “Locus gene repertoire per IMGT annotated assembly^17^”).

#### 4.1.2 Allelic polymorphism

The gorilla IG genes are shown to be polymorphic (Supplementary Table 1-3) as for other primate species, the different alleles resulting from nucleotide substitutions and/or nucleotide insertions or deletions which may lead to a modification of the gene functionality. For some alleles of genes, eg. IGHV7-40, IGHV7-40D, IGKV2-23 and IGKV2-38, the functionality was altered due to insertion of repeated foreign IG DNA sequences “Repeat regions”, mostly the LINEs and SINEs families. The proportion of these regions is over 60% in mammals (48).

The characterization of the allelic polymorphism was based on the IMGT unique numbering (34,35) for an easy comparison of codons and amino acids sequences of V, D, J and C regions or exons (34). The dynamic gene tables per IMGT group and per species^18^ lists the alleles of IG genes and their corresponding functionality. In the gene table, a scoring system based on one to three stars indicates that a given allele was identified in one, two or more genomic sequences. In the context of the evolution of high throughput sequencing technologies, more than one star would confirm the existence of the alleles and eliminate suspicion of sequencing errors. Following the analysis of the four assemblies 63% of IGH, 57% of IGL, 64% of IGK genes are shown to be polymorphic. The higher number of annotated assemblies in the future, the more accurate this estimation will be.

#### 4.1.3 Gene insertion/deletion and CNVs in IGHV cluster

The analysis of the IGH locus within the four assemblies of gorilla allowed us to report variations on the gorilla genome between individuals, in particular Copy Number Variations -CNVs-(Figure 5, Supplementary Table 1). Among these, some CNVs have already been described in human and some non-human primates (49), indicating that human CNVs could be no specific to the human species. Gazave and colleagues (50) observed that the majority of CNVs are not species specific, and they are consistent with species phylogenetic relationships. Shared CNVs might be the result of ancient structural polymorphism retention, as well as high segmental duplication activity, that facilitate recurrent loss or creation of new copies via Non Allelic Homologous Recombination (50).

**Figure 5.**
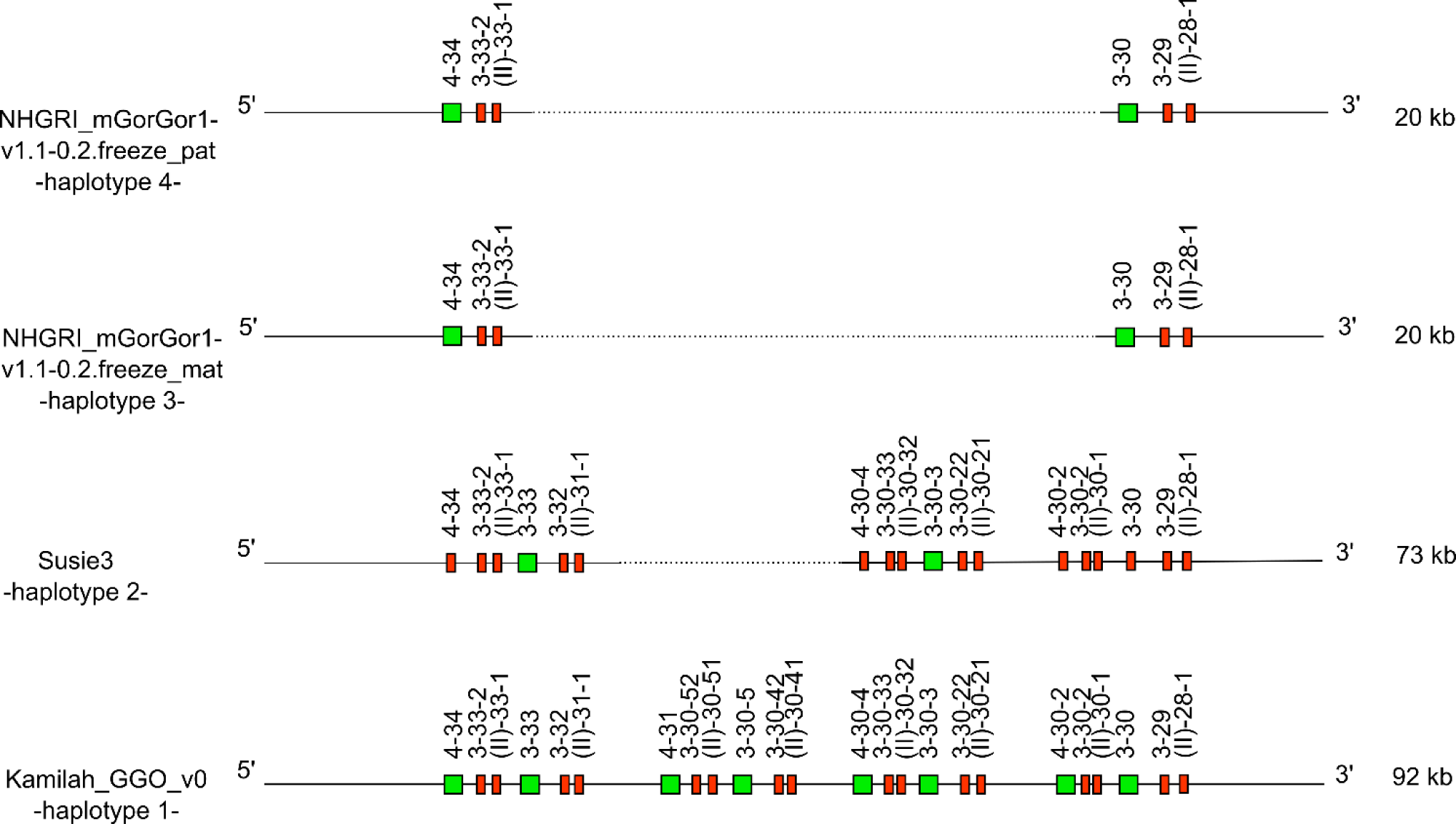
CNV3- of *Gorilla gorilla gorilla* IGH locus (from IGHV4-34 to IGHV(II)-28-1) which is shared with *Homo sapiens*. IGHV4-28 is absent from the gorilla genome. The first gorilla haplotype contains all genes of human counterpart CNV3 form C. The new CNV3 form H shows deletion of six genes, and the new CNV3 form I shows a deletion of 18 genes. Lengths mentioned on the right, are different depending on the insertion/deletion of genes.

According to the human and gorilla genes organization in IGH locus, this could be illustrated by the human CNV3 (30) (represented in the “Human (Homo sapiens) IGH CNV3 IMGT^19^” Web page).

The counterpart of this CNV was also identified in gorilla with the CNV3 form (gene content of the CNV between the 3’ and 5’ limit) C in Kamilah_GGO_v0 -haplotype 1- and two new CNV3 forms, not described in human, called forms H and I, apparently gorilla specific.

It is worth noting that additional human CNVs, namely CNV1, CNV6 and CNV7 could be identified in gorilla IGH loci (with variation in number of genes), with gorilla specific CNV forms (Supplementary Table 1).

New sets of IG genes have been identified in gorilla loci compared to human, five of them could be associated with new gorilla specific CNVs, called potential CNVp1 to CNVp5 (Supplementary Table 1). However, one other set composed of the six IGHD genes absent from Kamilah_GGO_v0 assembly because of 20 kb gap are not proposed as potential CNV.

### 4.2 IG constant genes and CNVs

#### 4.2.1 IGH C-GENE classes and subclasses characterization

The five IG classes (IgA, IgE, IgD, IgM and IgG) are characterized by different heavy chain constant regions, coded by the constant genes of the IGH locus (51). In this study, we highlighted a highly similar IGHC gene organization between gorilla and human (Supplementary Table 2, Supplementary Figures 1-4). This corroborates previous work describing gorilla, chimpanzees and human and orthologous genes (45). However, even if the presence of several IGHG were already mentioned (45,46), to our knowledge, this is the first study that identifies and characterizes the nine distinct IGHC genes and in particular the six IGHG genes. Indeed, the three IGHG3 genes IGHG3A, IGHG3B and IGHG3C are present in three of four assemblies. This would confirm that duplications continued to occur especially in the clade of IgG3 where gorillas and chimpanzees created an additional IgG, which reflects evolutionary instability in the locus. The gorilla IgG3 isotype is characterized by three constant domains and a variant number of hinge regions, from two to five (2-5 for IgG3A, 2 for IgG3B and 4 for IgG3C) (see Dynamic gene tables per IMGT group and per species). The hinge in IgG3 immunoglobulin class is therefore longer than the hinge regions of the other IgG subclasses (51), except for IGHGP. According to the number of hinge regions (and to high similarity of domain exons), the gorilla IGHG3C gene seems to be the most closely matched to the human IGHG3.

Sequences of IGHG2, IGHG4, IGHE and IGHA2 were not detected within the IGH loci of the Kamilah_GGO_v0 nor that of Susie3. In the latest assembly, we found the four genes on contig CYUI03001141.1^20^ (data not shown), associated to the related bioproject but not assembled on chromosome 14. These four genes were found and annotated within the IGH locus in both NHGRI_mGorGor1-v1.1-0.2.freeze_mat and NHGRI_mGorGor1-v1.1-0.2.freeze_pat haploid genomes.

Human, chimpanzees and gorillas seem to share a common ancestral duplication of the IGHG, IGHE and IGHA genes (52), that likely had taken place in their common ancestor. Therefore, the IGHE and IGHEP1 genes were linked to the IGHA2 and IGHA1 respectively in the gorilla genome (52). In order to confirm the correct gene name assignment of IGHEP1, IGHA1 and IGHE, IGHA2 genes, the characteristic length of IGHA genes was taken into account. On one hand, our results contradict those reported in (52), regarding the hinge region length of the gorilla IGHA1 compared to those of human and chimpanzee: we noticed the presence of duplication in the Hinge region of gorilla IGHA1 that also occurred in human and chimpanzee. We confirm that the Hinge of the third allele of gorilla is shorter because of the deletion of 2 nucleotides, leading to frameshift in the reading frame (Figure 6 (A)). On the other hand, we concur with the assumption made by the same scientific team that, the IGHA2 gene was derived from the prototype IGHA1, by the 15 bp deletion in the Hinge region, before its duplication, which seems to have occurred before divergence of the three species (human, chimpanzees and gorillas) (Figure 6 (B)).

**Figure 6.**
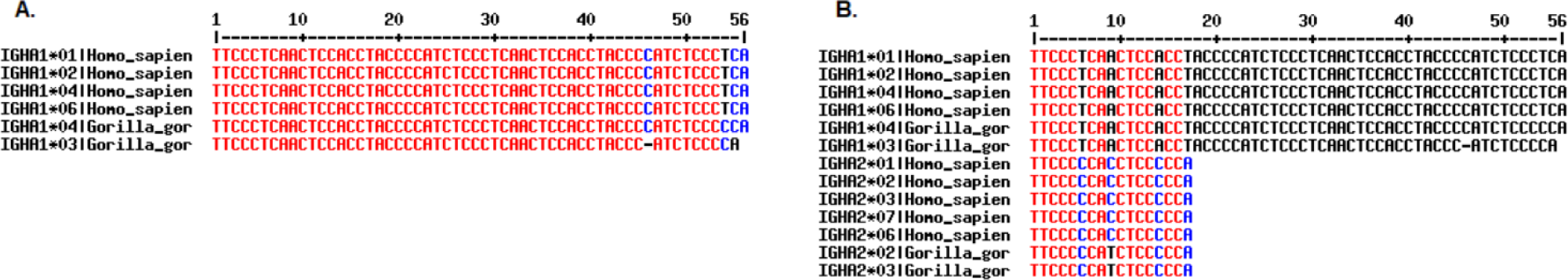
(A) Alignment of *Homo sapiens* and *Gorilla gorilla gorilla* IGHA1 allele sequences. (B) Alignment of *Homo sapiens* and *Gorilla gorilla gorilla* IGHA1 and IGHA2 alleles sequences. Alignments generated using MultAlin software (53).

Two IGHE genes were annotated on the main IGH locus of gorilla: IGHE, and IGHEP1 which is truncated in 5’. It seems that among the hominoids, only the gorilla and human genomes contained three IGHE genes (54), two in the main locus, and one outside. Note that the detected IGHE gene outside the main locus of gorilla (data not shown), is the human IGHEP2 counterpart (processed gene outside the main IGH locus in human genome).

It should be noted that IGHC genes from *Gorilla gorilla* species have been previously annotated and published on the IMGT site. However, because of unknown subspecies, and partial and/or non identical sequences they were not reassigned according to this study (the *Gorilla gorilla* IGHG3 could not be assigned to a subtype IGHG3A, IGHG3B, or IGHG3C, and new allele numbers were assigned to the genes from the four assemblies if the genes was already published in the “Gene table: Western gorilla (*Gorilla gorilla*) IGHC^21^”).

#### 4.2.2 IGLJ and IGLC genes

An additional J-C-CLUSTER was identified in the gorilla IGL locus of the NHGRI_mGorGor1-v1.1-0.2.freeze_pat assembly compared to available IMGT annotated human assemblies. The (55) study comparing IGL sequences between different human populations revealed that some human populations could have up to four additional IGLC genes, most likely linked to a junction gene, localized between the IGLC2 and IGLC3 genes (see “Locus representation: human (*Homo sapiens*) IGL^22^” on IMGT Web site). As found in the same location in the gorilla counterpart, this represents a form of CNV, with 99% and 100% identity between IGLC2 and IGLC2A, and between IGLJ2 and IGLJ2A, respectively.

### 4.3 Gorilla and human IGK loci analysis

The IGK locus has a reverse orientation on the chromosome 2A in three assemblies, but is forward on chromosome 2A of the Susie3 assembly: we noticed that this unexpected locus orientation was also observed for the dog, *Canis lupus familiaris*: the IGK locus is REV for the CanFam3.1 and FWD for the Basenji_breed-1.1, both assemblies annotated in IMGT (see “Locus in genome assembly: dog (Canis lupus familiaris) IGK locus^23^”).

Indeed, dog and gorilla assemblies were built using the comparison with the human one: as the human IGK locus is composed of a proximal IGK in REV orientation and a duplicated part, the distal IGK locus in FWD orientation on chromosome 2. As neither the dog nor the gorilla shows duplicated part in their IGK locus, this individual change in the IGK locus orientation could be linked to a methodology artefact.

The gorilla IGK locus contains six additional IGKV genes in the 5’ side of the V-CLUSTER with no identified human counterpart up to now.

Our results indicating the detection of IGK genes only between gorilla IGK 5’ and 3’ “IMGT bornes” confirm the existence of one and unique IGK locus. As cited in (56), no indication of a duplication within the IGK locus was obtained in establishing the *Pan troglodytes* and the *Gorilla gorilla* maps. Whereas the human IGK locus has two V-CLUSTER in inverse orientation to each other, which are very similar but not identical, called the proximal (p) and the distal (d) locus (57).

Our findings show that the genes of gorilla IGK locus present high percentage of identity with human genes of the distal IGK V-CLUSTER, and similar structural organization, especially since we found in gorilla three genes corresponding to the three additional human genes counterpart of the distal V-CLUSTER, that have no duplicate equivalent on the human proximal V-CLUSTER: IGKV6-41, IGKV1-42 and IGKV1-43 (Supplementary Table 2). The divergence between proximal and distal V-CLUSTER is largely due to points of mutations indicating that the duplication is an evolutionary (58) involved deletions in some regions on the proximal locus, that must have occurred after duplication of the locus (59,60). The absence of two V-CLUSTER in the IGK locus in chimpanzee and gorilla means that the duplication in human IGK locus occurred after the branch-point human and great ape evolution (56).

### 4.4 *Gorilla gorilla gorilla* chromosome nomenclature

Up to January 8, 2024, non-human primates chromosomes were named by homology with the ones of human. The common ancestor of gorillas, chimpanzees and human had 24 pairs of chromosomes (61). Great apes have conserved the same number of diploid chromosomes, whereas modern human possess 23 pairs (2n=46) due to a telomeric fusion of chromosomes 2A and 2B. Most chromosomes appear to be similar between the three species, with the remaining differences between chromosomes consisting of inversions of chromosome segments and variations in constitutive heterochromatin (61). Based on this nomenclature, we were able to localize the three IG loci on the same chromosomes as human: the IGH locus on chromosome 14, the IGK locus on chromosome 2A in the gorilla and the locus IGL on chromosome 22.

Since January 8, 2024, the non-human primate chromosome pairs were updated and renamed from 1 to 24. For gorilla, only assemblies of KB3781 individual have been updated (NHGRI_mGorGor1-v2.0_pri, NHGRI_mGorGor1-v2.0_mat, NHGRI_mGorGor1-v2.0_pat). Therefore, the IGH locus now resides on chromosome 15, the IGK locus on chromosome 12, and the IGL locus on chromosome 23. In the context of phylogenetic studies between apes and human, and gorilla individuals, the previous version of assemblies was more appropriate, in our opinion, in terms of the close phylogenetic relationship between gorilla and human and their common ancestor.

We strongly believe that this sort of important change should be taken after consultation of the scientific community and clear prior communication before implementation.

### 4.5 Assemblies of “Kamilah” individual

The Kamilah_GGO_v0, Kamilah_GGO_hifiasm-v0.15.2.pri and Kamilah_GGO_hifiasm-v0.15.2.alt assemblies (Figure 1) originate from the same individual, Kamilah. The analysis of Kamilah_GGO_v0 (oldest Kamilah assembly) IG loci is an integral part of this study, while the analysis of Kamilah_GGO_hifiasm-v0.15.2.pri and Kamilah_GGO_hifiasm-v0.15.2.alt, NCBI assemblies at “Contig” level were not integrated, because they do not correspond to our assembly selection criteria for this study.

Upon a preliminary analysis of the IG loci in the two most recent assemblies, similar gene organization within the same locus was confirmed as expected. However, we observed variations in gene number within the IGH and IGL loci but not in IGK locus (Figure 7).

**Figure 7.**
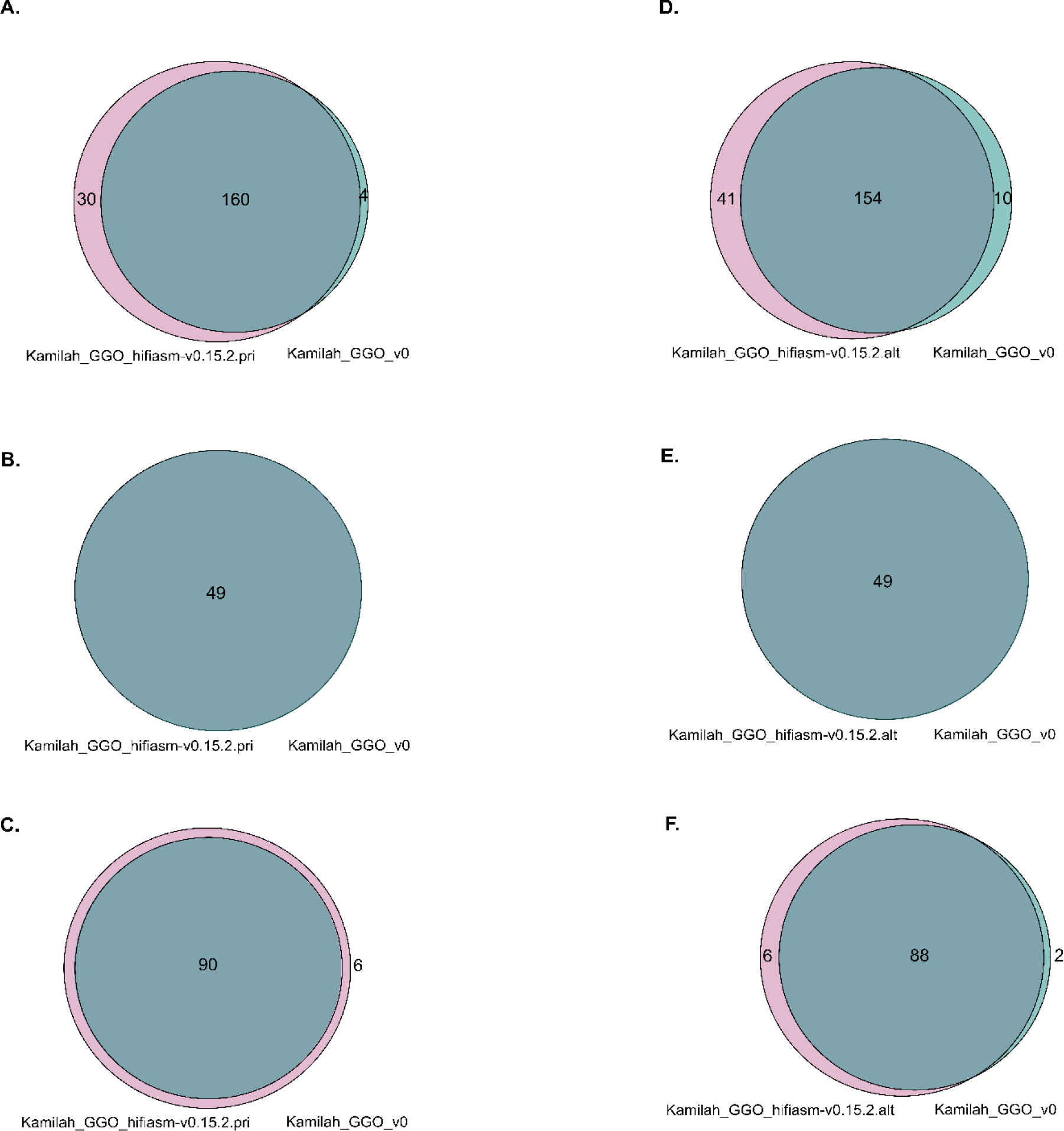
(A), (B) and (C) Venn diagrams represent shared and different IG sequences between Kamilah_GGO_hifiasm-v0.15.2.pri and Kamilah_GGO_v0 assemblies of the same individual, Kamilah. (D), (E) and (F) Venn diagrams represent shared and different IG sequences between Kamilah_GGO_hifiasm-v0.15.2.alt and Kamilah_GGO_v0 assemblies of Kamilah individual. The gray intersection indicates sequences that have 100% identity. The pink color represents new sequences in Kamilah_GGO_hifiasm-v0.15.2.pri / Kamilah_GGO_hifiasm-v0.15.2.alt compared to Kamilah_GGO_v0. The blue-green color indicates sequences that are only present in Kamilah_GGO_v0.

Considering the origin of the data, the source of differences could be linked to the biological material and/or to the sequencing technologies:

-Given that the Kamilah_GGO_v0 genome was obtained from a primary cultured fibroblast cell line, and both Kamilah_GGO_hifiasm-v0.15.2.pri and Kamilah_GGO_hifiasm-v0.15.2.alt were obtained from the cell line, the different tissue types influence genomic stability. On one hand, cell lines are frequently derived from a single cell, yet they might accrue genetic changes over time due to extended cultivation. Primary cells, on the other hand, may better reflect the individual’s genetic composition, although they may comprise a variety of cell types and are subject to culture changes.

-Depending on the sequencing and assembly methods, the identified sequence variations might be linked to methodological parameters such as sequencing coverage, read depth, mapping quality, assembly contiguity, and accuracy. Illumina technology produces shorter, lower-quality reads than PacBio technology, which produces longer reads (62). PacBio Sequel technology offers a higher consensus accuracy than PacBio RS II (63).

The Kamilah_GGO_hifiasm-v0.15.2.pri genome has more genes than the Kamilah_GGO_v0 genome. This is because Kamilah_GGO_v0 is missing 15 IGHD genes (due to a gap at this position) and nine IGHC genes in the IGH locus. Additionally, the Kamilah_GGO_hifiasm-v0.15.2.pri genome contains additional IGH and IGL V genes which are not present in Kamilah_GGO_v0.

However, several duplications (in V-CLUSTER and C-CLUSTER in IGH locus) seem to occur in the Kamilah_GGO_hifiasm-v0.15.2.alt assembly. These would be more likely the result of sequencing and/or assembly errors, although the Hifiasm assembly technique (utilized for the two recent assemblies of Kamilah) has a clear advantage over the other assemblers investigated in (64), including the FALCON assembly method, the one used for Kamilah_GGO_v0.

These preliminary studies allowed us “to fill the gaps” in the D-CLUSTER of IGH locus of Kamilah_GGO_v0, and to confirm the existence of nine additional IGHC genes in the Kamilah individual. However, an in-depth and complete analysis would be needed to interpret the meaning of differences between the three concerned assemblies.

## 5 Conclusion

Through the deciphering of the immunoglobulin genes at the three IG loci (IGH, IGK, and IGL), from four Western lowland gorilla (*Gorilla gorilla gorilla*) NCBI genome assemblies, IMGT^®^ provides a consistent overview of the organization and description of these loci and the potential individual variations in this closely related primate to human.

Due to the highly similar organization of gorilla and human loci and the high percentage of identity between IG genes in gorilla and human, the IMGT names of gorilla IG genes were mostly assigned according to their human counterparts.

The IG loci and the gene characterization, thanks to IMGT gene nomenclature and IMGT standards, highlighted characteristics of the gorilla genome:

> As in human, the gorilla IGH locus shows the greatest variability between individuals in terms of gene content. Several known human CNVs were identified in the gorilla IGH locus, along with new forms, as well as other potentially new CNVs called CNVp until their confirmation in other assemblies.
>
> The analysis of the organization of IG constant genes in the IGH locus from several individuals helped to better estimate the number of IGHG genes, which had been previously underestimated based on a single assembly (46), particularly with the characterization of three IGHG3 genes.
>
> The IGK locus is remarkably homogeneous in the four assemblies: it is characterized by the absence of IGKV locus duplication, which occurred in the human IGK locus, in addition, the IGKV gene cluster seems to be closer to the distal human V locus.
>
> The IGL locus comprises a CNV in the J-C-CLUSTER, which was suspected in the human IGL locus but had not been shown in the IMGT-annotated IGL locus until now.

The analysis of these loci generated a large amount of expertly curated data from the three gorilla individuals, which are distributed through the IMGT website resources, databases, tools, and web resources compiled in Supplementary Table 4. Although data from three individuals cannot reflect those of an entire population, they have enriched our immunogenetics knowledge of this species, a closely related primate to human, and will continue to evolve with the publication and expertise of new genome assemblies based on improved sequencing technologies and data from an increasing number of individuals.

The analysis of immunogenetic data is crucial in current immunology research. Studying great apes like gorillas, which are central to the Hominoidea group, offers valuable insights into primate evolution.

## 7 Author Contributions

CD conceived, analyzed the data, discussed and drafted the manuscript. GF and JJM conceived, analyzed the data and drafted the manuscript. VG conceived, discussed and drafted the manuscript, and supervised the write up of this work. SK conceived and supervised the findings and the writing of this manuscript.

## 9 Acknowledgments

We thank all members of the IMGT^®^ team for their expertise and constant motivation. We recognize the groundbreaking contributions of Marie-Paule Lefranc, the founder of IMGT^®^. IMGT^®^ is member of the French Infrastructure Institut Français de Bioinformatique (IFB) as well as member of BioCampus, MAbImprove and IBiSA. This work was granted access to the High Performance Computing (HPC) resources of Meso@LR and of Centre Informatique National de l’Enseignement Supérieur (CINES), to Très Grand Centre de Calcul (TGCC) of the Commissariat à l’Energie Atomique et aux Energies Alternatives (CEA) and to Institut du développement et des ressources en informatique scientifique (IDRIS) under the allocation 036029 (2010-2024) made by GENCI (Grand Equipement National de Calcul Intensif).

## 10 Supplementary Material

The supplementary Material for this article can be found online.

## 12 Data Availability Statement

The datasets generated for this study can be found in the online IMGT^®^ databases, tools and Web resources (https://www.imgt.org/), and in the online Third Party Annotation (TPA) (https://www.ncbi.nlm.nih.gov/genbank/tpa/).

## Footnotes

1 https://www.imgt.org/

2 https://www.imgt.org/vquest/refseqh.html#refdir2

3 https://www.imgt.org/IMGTScientificChart/SequenceDescription/IMGTassemblyselection.html

4 https://www.ncbi.nlm.nih.gov/datasets/docs/v2/glossary/

5 https://www.imgt.org/IMGTindex/IMGTborne.php

6 https://www.imgt.org/IMGTScientificChart/SequenceDescription/IMGTfunctionality.html

7 https://www.imgt.org/IMGTScientificChart/Nomenclature/IMGTnomenclature.html

8 https://www.imgt.org/IMGTrepertoire/

9 https://www.ncbi.nlm.nih.gov/genbank/tpa/

10 https://www.imgt.org/IMGTindex/labels.php

11 https://www.imgt.org/IMGTindex/IGHVclans.php

12 https://www.imgt.org/IMGTrepertoire/LocusGenes/locusdesc/assembly_compare.php

13 https://www.imgt.org/IMGTrepertoire/LocusGenes/genetable/autotable.php

14 https://www.imgt.org/IMGTrepertoire/LocusGenes/bornes/bornesIGK.html

15 https://www.imgt.org/IMGTrepertoire/LocusGenes/bornes/bornesIGL.html

16 https://www.imgt.org/IMGTindex/IGLVclans.php

17 https://www.imgt.org/IMGTrepertoire/LocusGenes/locusdesc/assembly_compare.php

18 https://www.imgt.org/IMGTrepertoire/LocusGenes/genetable/autotable.php

19 https://www.imgt.org/IMGTrepertoire/LocusGenes/locus/human/IGH/CNV/Hu_IGHCNV3.html

20 https://www.ncbi.nlm.nih.gov/nuccore/CYUI03001141.1

21 https://www.imgt.org/IMGTrepertoire/index.php?section=LocusGenes&repertoire=genetable&species=Gorilla&group=I <u>GHC</u>

22 https://www.imgt.org/IMGTrepertoire/index.php?section=LocusGenes&repertoire=locus&species=human&group=IGL

23 https://www.imgt.org/IMGTrepertoire/index.php?section=LocusGenes&repertoire=locusAssembly&species=dog&group=IGK

## 13 References

1. Lefranc MP, Lefranc G. The Immunoglobulin FactsBook. Academic Press; 2001. 472 p.

2. Lefranc MP, Lefranc G. The T Cell Receptor FactsBook. Elsevier; 2001. 413 p.

3. Marshall JS, Warrington R, Watson W, Kim HL. An introduction to immunology and immunopathology. Allergy Asthma Clin Immunol. sept 2018;14(S2):49.

4. Tonegawa S. Somatic generation of antibody diversity. Nature. avr 1983;302(5909):575-81.

5. Giudicelli V, Lefranc MP. Ontology for immunogenetics: the IMGT-ONTOLOGY. Bioinformatics. 1 déc 1999;15(12):1047-54.

6. Das S, Nozawa M, Klein J, Nei M. Evolutionary dynamics of the immunoglobulin heavy chain variable region genes in vertebrates. Immunogenetics. janv 2008;60(1):47-55.

7. Hirano M, Das S, Guo P, Cooper MD. The Evolution of Adaptive Immunity in Vertebrates. In: Advances in Immunology [Internet]. Elsevier; 2011 [cité 25 juin 2024]. p. 125-57. Disponible sur: https://linkinghub.elsevier.com/retrieve/pii/B9780123876645000042

8. Tanaka T, Nei M. Positive darwinian selection observed at the variable-region genes of immunoglobulins. Mol Biol Evol. sept 1989;6(5):447-59.

9. Lefranc MP, Giudicelli V, Duroux P, Jabado-Michaloud J, Folch G, Aouinti S, et al. IMGT®, the international ImMunoGeneTics information system® 25 years on. Nucleic Acids Research. 28 janv 2015;43(D1):D413-22.

10. Manso T, Folch G, Giudicelli V, Jabado-Michaloud J, Kushwaha A, Nguefack Ngoune V, et al. IMGT® databases, related tools and web resources through three main axes of research and development. Nucleic Acids Research. 7 janv 2022;50(D1):D1262-72.

11. Lefranc MP, Giudicelli V, Ginestoux C, Jabado-Michaloud J, Folch G, Bellahcene F, et al. IMGT(R), the international ImMunoGeneTics information system(R). Nucleic Acids Research. 1 janv 2009;37(Database):D1006-12.

12. Lefranc MP. Immunoglobulin and T Cell Receptor Genes: IMGT® and the Birth and Rise of Immunoinformatics. Front Immunol [Internet]. 2014 [cité 25 juin 2024];5. Disponible sur: http://journal.frontiersin.org/article/10.3389/fimmu.2014.00022/abstract

13. Grow DA, McCarrey JR, Navara CS. Advantages of nonhuman primates as preclinical models for evaluating stem cell-based therapies for Parkinson’s disease. Stem Cell Research. sept 2016;17(2):352-66.

14. Bailey JA, Eichler EE. Erratum: Primate segmental duplications: crucibles of evolution, diversity and disease. Nat Rev Genet. nov 2006;7(11):898-898.

15. Symmons O, Varadi A, Aranyi T. How Segmental Duplications Shape Our Genome: Recent Evolution of ABCC6 and PKD1 Mendelian Disease Genes. Molecular Biology and Evolution. 5 août 2008;25(12):2601-13.

16. Scally A, Dutheil JY, Hillier LW, Jordan GE, Goodhead I, Herrero J, et al. Insights into hominid evolution from the gorilla genome sequence. Nature. mars 2012;483(7388):169-75.

17. Rogers J, Gibbs RA. Comparative primate genomics: emerging patterns of genome content and dynamics. Nat Rev Genet. mai 2014;15(5):347-59.

18. Nguefack Ngoune V, Bertignac M, Georga M, Papadaki A, Albani A, Folch G, et al. IMGT® Biocuration and Analysis of the Rhesus Monkey IG Loci. Vaccines. mars 2022;10(3):394.

19. Wilming LG, Hart EA, Coggill PC, Horton R, Gilbert JGR, Clee C, et al. Sequencing and comparative analysis of the gorilla MHC genomic sequence. Database [Internet]. 1 janv 2013 [cité 25 juin 2024];2013. Disponible sur: https://academic.oup.com/database/article/doi/10.1093/database/bat011/330685

20. Van Loghem E, De Lange G. Immunoglobulin Epitopes in Primates. Vox Sanguinis. déc 1979;37(6):329-37.

21. Pégorier P, Bertignac M, Chentli I, Nguefack Ngoune V, Folch G, Jabado-Michaloud J, et al. IMGT® Biocuration and Comparative Study of the T Cell Receptor Beta Locus of Veterinary Species Based on Homo sapiens TRB. Front Immunol [Internet]. 5 mai 2020 [cité 25 juin 2024];11. Disponible sur: https://www.frontiersin.org/journals/immunology/articles/10.3389/fimmu.2020.00821/full

22. Kitts PA, Church DM, Thibaud-Nissen F, Choi J, Hem V, Sapojnikov V, et al. Assembly: a resource for assembled genomes at NCBI. Nucleic Acids Res. 4 janv 2016;44(D1):D73-80.

23. NCBI Resource Coordinators. Database Resources of the National Center for Biotechnology Information. Nucleic Acids Res. 4 janv 2017;45(D1):D12-7.

24. Finstermeier K, Zinner D, Brameier M, Meyer M, Kreuz E, Hofreiter M, et al. A Mitogenomic Phylogeny of Living Primates. Stanyon R, éditeur. PLoS ONE. 16 juill 2013;8(7):e69504.

25. Gordon D, Huddleston J, Chaisson MJP, Hill CM, Kronenberg ZN, Munson KM, et al. Long-read sequence assembly of the gorilla genome. Science. 1 avr 2016;352(6281):aae0344.

26. O’Leary NA, Cox E, Holmes JB, Anderson WR, Falk R, Hem V, et al. Exploring and retrieving sequence and metadata for species across the tree of life with NCBI Datasets. Sci Data. 5 juill 2024;11:732.

27. Xu X, Arnason U. A complete sequence of the mitochondrial genome of the western lowland gorilla. Molecular Biology and Evolution. 1 mai 1996;13(5):691-8.

28. Mao Y, Harvey WT, Porubsky D, Munson KM, Hoekzema K, Lewis AP, et al. Structurally divergent and recurrently mutated regions of primate genomes. Cell. mars 2024;187(6):1547–1562.e13.

29. Altschul SF, Gish W, Miller W, Myers EW, Lipman DJ. Basic local alignment search tool. Journal of Molecular Biology. oct 1990;215(3):403-10.

30. Lefranc MP, Lefranc G. IMGT®Homo sapiens IG and TR Loci, Gene Order, CNV and Haplotypes: New Concepts as a Paradigm for Jawed Vertebrates Genome Assemblies. Biomolecules. mars 2022;12(3):381.

31. Giudicelli V. IMGT/LIGM-DB, the IMGT(R) comprehensive database of immunoglobulin and T cell receptor nucleotide sequences. Nucleic Acids Research. 1 janv 2006;34(90001):D781-4.

32. Lane J, Duroux P, Lefranc MP. From IMGT-ONTOLOGY to IMGT/LIGMotif: the IMGT® standardized approach for immunoglobulin and T cell receptor gene identification and description in large genomic sequences. BMC Bioinformatics. déc 2010;11(1):223.

33. Sievers F, Higgins DG. Clustal Omega for making accurate alignments of many protein sequences. Protein Science. janv 2018;27(1):135-45.

34. Lefranc MP. IMGT Unique Numbering for the Variable (V), Constant (C), and Groove (G) Domains of IG, TR, MH, IgSF, and MhSF. Cold Spring Harb Protoc. juin 2011;2011(6):pdb.ip85.

35. Lefranc MP, Pommié C, Ruiz M, Giudicelli V, Foulquier E, Truong L, et al. IMGT unique numbering for immunoglobulin and T cell receptor variable domains and Ig superfamily V-like domains. Developmental & Comparative Immunology. janv 2003;27(1):55-77.

36. Giudicelli V, Lefranc MP. IMGT-ONTOLOGY 2012. Front Gene [Internet]. 2012 [cité 25 juin 2024];3. Disponible sur: http://journal.frontiersin.org/article/10.3389/fgene.2012.00079/abstract

37. Wong K. Scientific American. 2014 [cité 24 juin 2024]. Tiny Genetic Differences between Humans and Other Primates Pervade the Genome. Disponible sur: https://www.scientificamerican.com/article/tiny-genetic-differences-between-humans-and-other-primates-pervade-the-genome/

38. Giudicelli V. IMGT/GENE-DB: a comprehensive database for human and mouse immunoglobulin and T cell receptor genes. Nucleic Acids Research. 17 déc 2004;33(Database issue):D256-61.

39. Cochrane G, Bates K, Apweiler R, Tateno Y, Mashima J, Kosuge T, et al. Evidence Standards in Experimental and Inferential INSDC Third Party Annotation Data. OMICS: A Journal of Integrative Biology. juin 2006;10(2):105-13.

40. Lefranc MP. From IMGT-ONTOLOGY DESCRIPTION Axiom to IMGT Standardized Labels: For Immunoglobulin (IG) and T Cell Receptor (TR) Sequences and Structures. Cold Spring Harb Protoc. juin 2011;2011(6):pdb.ip83.

41. Lemoine F, Correia D, Lefort V, Doppelt-Azeroual O, Mareuil F, Cohen-Boulakia S, et al. NGPhylogeny.fr: new generation phylogenetic services for non-specialists. Nucleic Acids Research. 2 juill 2019;47(W1):W260-5.

42. Letunic I, Bork P. Interactive Tree of Life (iTOL) v6: recent updates to the phylogenetic tree display and annotation tool. Nucleic Acids Research. 13 avr 2024;gkae268.

43. Levinson AI, Kozlowski L, Zheng Y, Wheatley L. B-cell superantigens: definition and potential impact on the immune response. J Clin Immunol. nov 1995;15(6 Suppl):26S-36S.

44. Deacy AM, Gan SKE, Derrick JP. Superantigen Recognition and Interactions: Functions, Mechanisms and Applications. Front Immunol. 20 sept 2021;12:731845.

45. Olivieri DN, Gambón Deza F. Immunoglobulin genes in Primates. Molecular Immunology. sept 2018;101:353-63.

46. Garzón-Ospina D, Buitrago SP. Immunoglobulin heavy constant gamma gene evolution is modulated by both the divergent and birth-and-death evolutionary models. Primates. nov 2022;63(6):611-25.

47. Barbié V, Lefranc MP. The Human Immunoglobulin Kappa Variable (IGKV) Genes and Joining (IGKJ) Segments. Exp Clin Immunogenet. 1998;15(3):171-83.

48. Wen-Hsiung LL, Dan G. Fundamentals of Molecular Evolution. First Edition. Vol. 252.

49. Watson CT, Steinberg KM, Huddleston J, Warren RL, Malig M, Schein J, et al. Complete Haplotype Sequence of the Human Immunoglobulin Heavy-Chain Variable, Diversity, and Joining Genes and Characterization of Allelic and Copy-Number Variation. The American Journal of Human Genetics. avr 2013;92(4):530-46.

50. Gazave E, Darré F, Morcillo-Suarez C, Petit-Marty N, Carreño A, Marigorta UM, et al. Copy number variation analysis in the great apes reveals species-specific patterns of structural variation. Genome Res. oct 2011;21(10):1626-39.

51. Attanasio R, Jayashankar L, Engleman CN, Scinicariello F. Baboon immunoglobulin constant region heavy chains: identification of four IGHG genes. Immunogenetics. nov 2002;54(8):556-61.

52. Ueda S, Matsuda F, Honjo T. Multiple recombinational events in primate immunoglobulin epsilon and alpha genes suggest closer relationship of humans to chimpanzees than to gorillas. J Mol Evol. mars 1988;27(1):77-83.

53. Corpet F. Multiple sequence alignment with hierarchical clustering. Nucleic Acids Research. 25 nov 1988;16(22):10881-90.

54. Ueda S, Takenaka O, Honjo T. A truncated immunoglobulin epsilon pseudogene is found in gorilla and man but not in chimpanzee. Proc Natl Acad Sci USA. juin 1985;82(11):3712-5.

55. Taub RA, Hollis GF, Hieter PA, Korsmeyer S, Waldmann TA, Leder P. Variable amplification of immunoglobulin λ light-chain genes in human populations. Nature. juill 1983;304(5922):172-4.

56. Ermert K, Mitlöhner H, Schempp W, Zachau HG. The immunoglobulin κ locus of primates. Genomics. févr 1995;25(3):623-9.

57. Brensing-Küppers J, Zocher I, Thiebe R, Zachau HG. The human immunoglobulin κ locus on yeast artificial chromosomes (YACs). Gene. juin 1997;191(2):173-81.

58. Schäble KF, Zachau HG. The variable genes of the human immunoglobulin kappa locus. Biol Chem Hoppe Seyler. nov 1993;374(11):1001-22.

59. Lautner-Rieske A, Huber C, Meindl A, Pargent W, Schäble KF, Thiebe R, et al. The human immunoglobulin x locus. Characterization of the duplicated A regions. Eur J Immunol. avr 1992;22(4):1023-9.

60. Huber C, Huber E, Lautner-Rieske A, Schäble KF, Zachau HG. The human immunoglobulin χ locus. Characterization of the partially duplicated L regions. Eur J Immunol. nov 1993;23(11):2860-7.

61. Yunis JJ, Prakash O. The Origin of Man: A Chromosomal Pictorial Legacy. Science. 19 mars 1982;215(4539):1525-30.

62. Rayamajhi N, Cheng CHC, Catchen JM. Evaluating Illumina-, Nanopore-, and PacBio-based genome assembly strategies with the bald notothen, *Trematomus borchgrevinki*. Rokas A, éditeur. G3 Genes|Genomes|Genetics. 4 nov 2022;12(11):jkac192.

63. Tvedte ES, Gasser M, Sparklin BC, Michalski J, Hjelmen CE, Johnston JS, et al. Comparison of long-read sequencing technologies in interrogating bacteria and fly genomes. Baltrus D, éditeur. G3 Genes|Genomes|Genetics. 17 juin 2021;11(6):jkab083.

64. Yu W, Luo H, Yang J, Zhang S, Jiang H, Zhao X, et al. Comprehensive assessment of 11 de novo HiFi assemblers on complex eukaryotic genomes and metagenomes. Genome Res. févr 2024;34(2):326-40.

