## Supplemental Tables 5 6 7 for "Deciphering of *Gorilla gorilla gorilla* Immunoglobulin Loci in Multiple Genome Assemblies and Enrichment of IMGT Resources"

**Supplementary table 5 numbers of Western lowland gorilla (*Gorilla gorilla gorilla*) functional F, Open reading frame O, pseudogene P IGHV gene per subgroup in IMGT annotated assemblies**

| IMGT Subgroup | Kamilah_GGO_v0 | | Susie3 | | NHGRI_mGorGor1-v1.1-0.2.freeze_mat | | NHGRI_mGorGor1-v1.1-0.2.freeze_pat | |
|---|---|---|---|---|---|---|---|---|
| | **Gene nb** | **Total** | **Gene nb** | **Total** | **Gene nb** | **Total** | **Gene nb** | **Total** |
| IGHV1 | 6 F + 1 O + 6 P | 13 | 4 F + 1 O + 10 P | 15 | 6 F + 1 O + 8 P | 15 | 6 F + 1 O + 6 P | 13 |
| IGHV2 | 2 F + 1 O + 1 P | 4 | 1 F + 1 O + 3 P | 5 | 2 F + 1 O + 1 P | 4 | 2 F + 1 O + 1 P | 4 |
| IGHV3 | 26 F + 26 P | 52 | 14 F + 2 O + 33 P | 49 | 21 F + 2 O + 20 P | 43 | 23 F + 1 O + 23 P | 47 |
| IGHV4 | 10 F + 1 P | 11 | 6 F + 1 O + 4 P | 11 | 7 F | 7 | 8 F | 8 |
| IGHV5 | 1 O + 1 P | 2 | 1 O + 1 P | 2 | 1 O + 1 P | 2 | 1 O + 1 P | 2 |
| IGHV6 | 1 F | 1 | 1 F | 1 | 1 F | 1 | 1 F | 1 |
| IGHV7 | 1 F + 1 O + 4 P | 6 | 1 F + 1 O + 3 P | 5 | 2 F + 1 O + 3 P | 6 | 1 F + 1 O + 3 P | 5 |
| IGHV8 | 1 P | 1 | 1 P | 1 | 1 P | 1 | 1 P | 1 |
| IGHV(II) | 29 P | 29 | 29 P | 29 | 24 P | 24 | 26 P | 26 |
| IGHV(III) | 14 P | 14 | 16 P | 16 | 16 P | 16 | 15 P | 15 |
| IGHV(IV) | 1 P | 1 | 1 P | 1 | 1 P | 1 | 1 P | 1 |
| Total number of IMGT genes | 46 F + 4 O + 84 P | 134 | 27 F + 7 O + 101 P | 135 | 39 F + 6 O + 75 P | 120 | 41 F + 5 O + 77 P | 123 |

**Supplementary table 6 numbers of Western lowland gorilla (*Gorilla gorilla gorilla*) functional F, Open reading frame O, pseudogene P IGKV gene per subgroup in IMGT annotated assemblies**

| IMGT Subgroup | Kamilah_GGO_v0 | | Susie3 | | NHGRI_mGorGor1-v1.1-0.2.freeze_mat | | NHGRI_mGorGor1-v1.1-0.2.freeze_pat | |
|---|---|---|---|---|---|---|---|---|
| | **Gene nb** | **Total** | **Gene nb** | **Total** | **Gene nb** | **Total** | **Gene nb** | **Total** |
| IGKV1 | 12 F + 1 O + 3 P | 16 | 10 F + 6 P | 16 | 13 F + 1 O + 3 P | 17 | 12 F + 1 O + 3 P | 16 |
| IGKV2 | 3 F + 1 O + 11 P | 15 | 2 F + 1 O + 10 P | 13 | 4 F + 1 O + 10 P | 15 | 4 F + 1 O + 10 P | 15 |
| IGKV3 | 4 F + 3 P | 7 | 4 F + 3 P | 7 | 4 F + 3 P | 7 | 4 F + 3 P | 7 |
| IGKV4 | 1 F | 1 | 1 P | 1 | 1 F | 1 | 1 F | 1 |
| IGKV5 | 1 F | 1 | 1 F | 1 | 1 F | 1 | 1 F | 1 |
| IGKV6 | 1 F + 1 O | 2 | 1 F + 1 O | 2 | 1 F + 1 O | 2 | 1 F + 1 O | 2 |
| IGKV7 | 1 F | 1 | 1 F | 1 | 1 F | 1 | 1 P | 1 |
| Total number of IMGT genes | 23 F + 3 O + 17 P | 43 | 19 F + 2 O + 20 P | 41 | 25 F + 3 O + 16 P | 44 | 23 F + 3 O + 17 P | 43 |

**Supplementary table 7 numbers of Western lowland gorilla (*Gorilla gorilla gorilla*) functional F, Open reading frame O, pseudogene P IGLV gene per subgroup in IMGT annotated assemblies**

| IMGT Subgroup | Kamilah_GGO_v0 | | Susie3 | | NHGRI_mGorGor1-v1.1-0.2.freeze_mat | | NHGRI_mGorGor1-v1.1-0.2.freeze_pat | |
|---|---|---|---|---|---|---|---|---|
| | Gene nb | Total | Gene nb | Total | Gene nb | Total | Gene nb | Total |
| IGLV1 | 3 F + 1 O + 2 P | 6 | 2 F + 1 O + 3 P | 6 | 3 F + 1 O + 2 P | 6 | 3 F + 3 P | 6 |
| IGLV2 | 4 F + 1 O + 2 P | 7 | 6 F + 1 O + 3 P | 10 | 5 F + 1 O + 3 P | 9 | 6 F + 1 O + 3 P | 10 |
| IGLV3 | 8 F + 2 O + 10 P | 20 | 11 F + 2 O + 11 P | 24 | 8 F + 2 O + 10 P | 20 | 11 F + 2 O + 11 P | 24 |
| IGLV4 | 1 F + 1 O + 3 P | 5 | 1 F + 1 O + 3 P | 5 | 1 F + 1 O + 3 P | 5 | 1 F + 1 O + 3 P | 5 |
| IGLV5 | 2 F + 1 O | 3 | 1 F + 2 P | 3 | 2 F + 1 O | 3 | 2 F + 1 P | 3 |
| IGLV6 | 1 F | 1 | 1 F | 1 | 1 F | 1 | 1 F | 1 |
| IGLV7 | 1 F + 1 O + 1 P | 3 | 1 F + 1 O + 1 P | 3 | 1 F + 1 O + 1 P | 3 | 1 F + 1 O + 1 P | 3 |
| IGLV8 | 1 F | 1 | 1 F | 1 | 1 F | 1 | 1 F | 1 |
| IGLV9 | 1 F | 1 | 1 F | 1 | 1 F | 1 | 1 F | 1 |
| IGLV10 | 1 O + 1 P | 2 | 1 O + 1 P | 2 | 1 O + 1 P | 2 | 1 O + 1 P | 2 |
| IGLV11 | 1 O | 1 | 1 O | 1 | 1 O | 1 | 1 O | 1 |
| IGLV(I) | 10 P | 10 | 11 P | 11 | 11 P | 11 | 11 P | 11 |
| IGLV(II) | 3 P | 3 | 4 P | 4 | 3 P | 3 | 4 P | 4 |
| IGLV(III) | 2 P | 2 | 2 P | 2 | 2 P | 2 | 2 P | 2 |
| IGLV(IV) | 4 P | 4 | 4 P | 4 | 4 P | 4 | 4 P | 4 |
| IGLV(V) | 2 P | 2 | 2 P | 2 | 2 P | 2 | 2 P | 2 |
| IGLV(VI) | 3 P | 3 | 4 P | 4 | 3 P | 3 | 4 P | 4 |
| IGLV(VII) | 2 P | 2 | 2 P | 2 | 2 P | 2 | 2 P | 2 |
| Total number of IMGT genes | 22 F + 9 O + 45 P | 76 | 25 F + 8 O + 53 P | 86 | 23 F + 9 O + 47 P | 79 | 27 F + 7 O + 52 P | 86 |
