## Supplemental Figures 1 2 3 4 for "Deciphering of *Gorilla gorilla gorilla* Immunoglobulin Loci in Multiple Genome Assemblies and Enrichment of IMGT Resources"

### Legend

V-GENE functionnal
V-GENE ORF
V-GENE pseudogene
J-GENE functionnal
J-GENE ORF
J-GENE pseudogene
D-GENE functionnal
D-GENE ORF
D-GENE pseudogene
C-GENE functionnal
C-GENE pseudogene
RPI  pseudogene

### Supp figure 1: Western lowland gorilla (*Gorilla gorilla gorilla*) IGH locus on chromosome 14 assembly Kamilah_GGO_v0

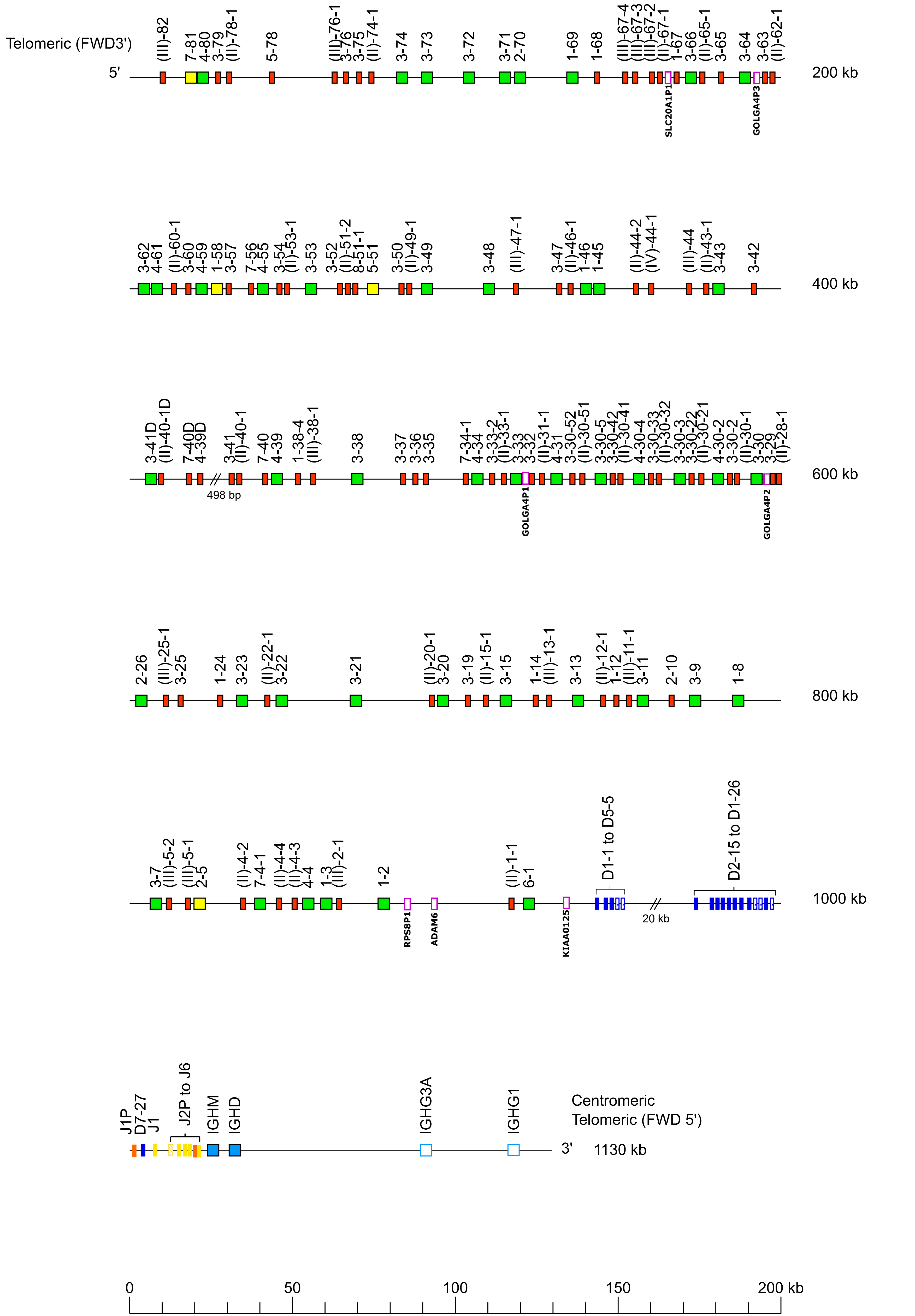

#### Supp figure 2: Western lowland gorilla (*Gorilla gorilla gorilla*) IGH locus on chromosome 14 assembly Susie3

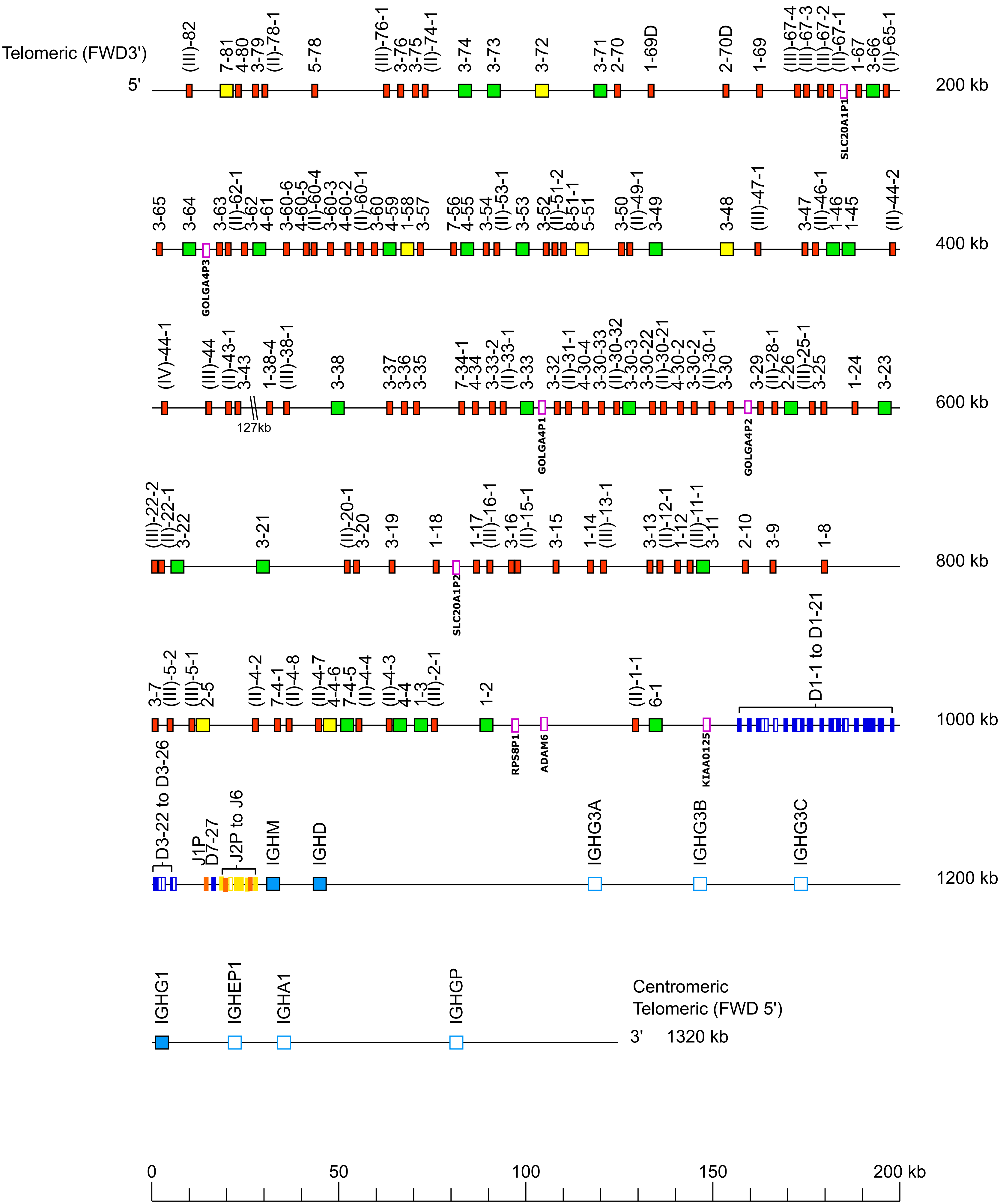

**Supp figure 3: Western lowland gorilla (*Gorilla gorilla gorilla*) IGH locus on chromosome 14 assembly NHGRI_mGorGor1-v1.1-0.2.freeze_mat**

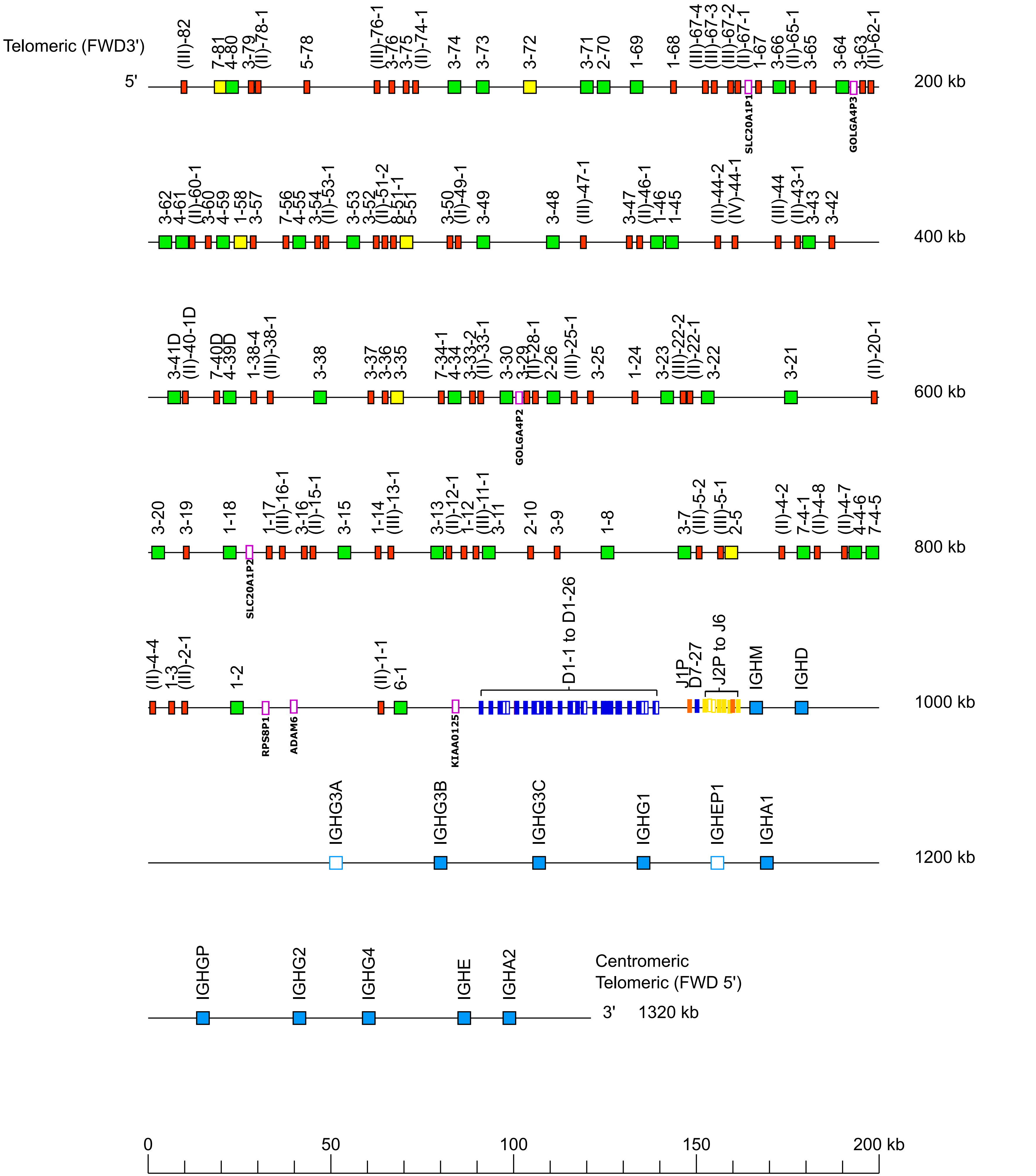

**Supp figure 4: Western lowland gorilla (*Gorilla gorilla gorilla*) IGH locus on chromosome 14 assembly NHGRI_mGorGor1-v1.1-0.2.freeze_pat**

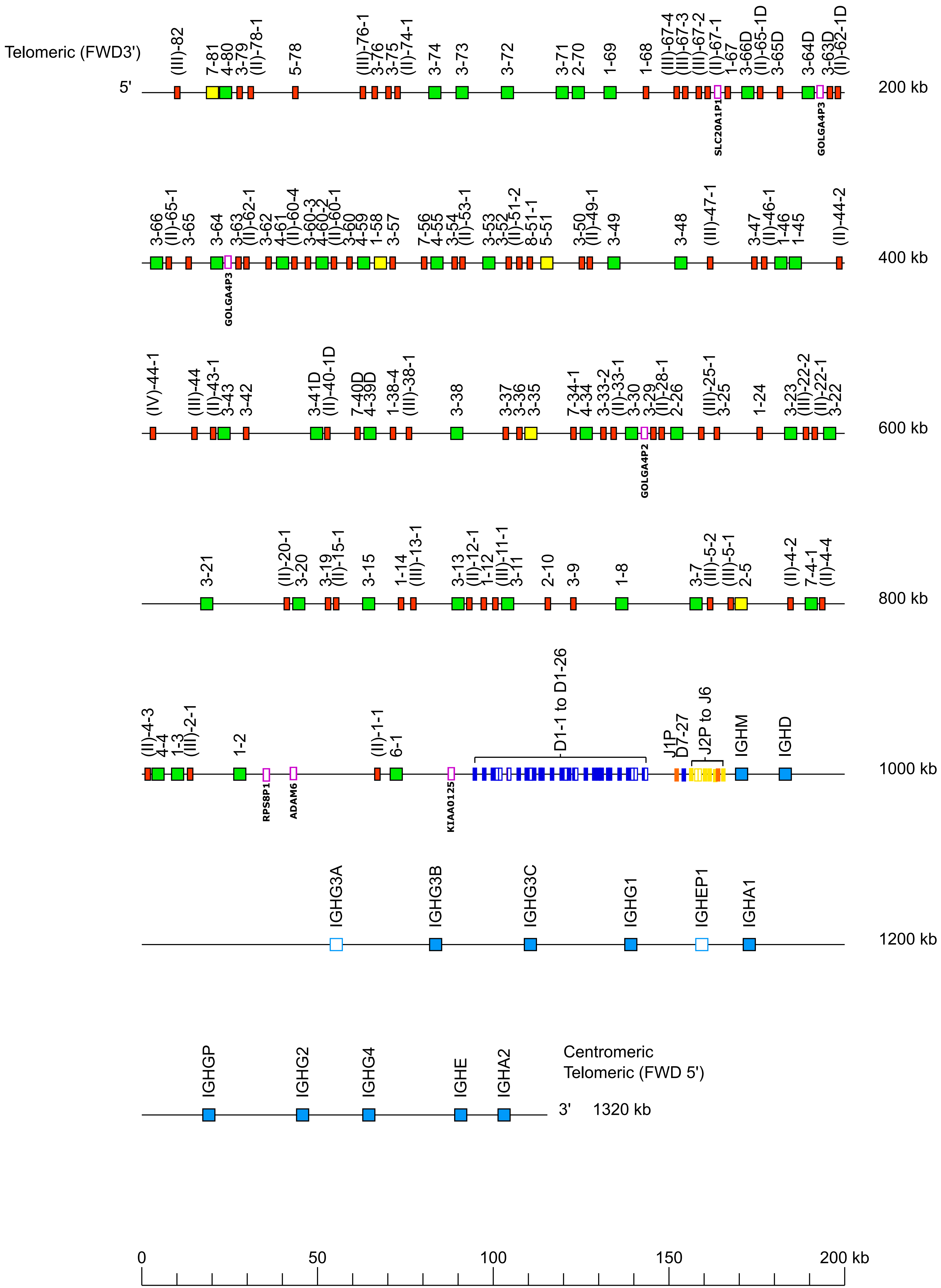
