## Supplemental Figures 5 6 7 8 for "Deciphering of *Gorilla gorilla gorilla* Immunoglobulin Loci in Multiple Genome Assemblies and Enrichment of IMGT Resources"

### Legend

- V-GENE functionnal
- V-GENE ORF
- V-GENE pseudogene
- J-GENE functionnal
- C-GENE functionnal
- 5' & 3' IMGT_borne gene functionnal

**Supp figure 5: Western lowland gorilla (*Gorilla gorilla gorilla*) IGK locus on chromosome 2A assembly Kamilah_GGO_v0**

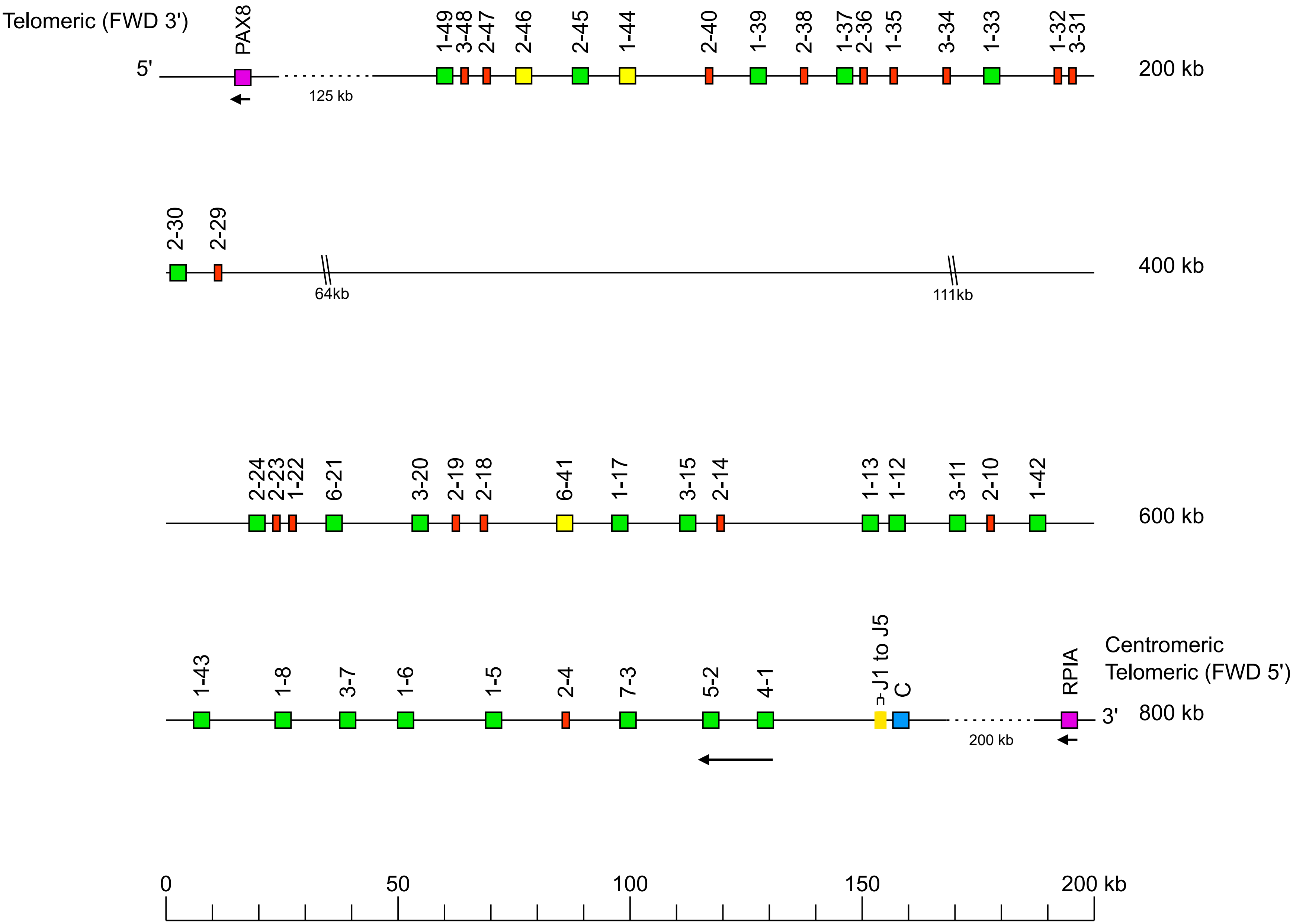

### Supp figure 6: Western lowland gorilla (*Gorilla gorilla gorilla*) IGK locus on chromosome 2A assembly Susie3

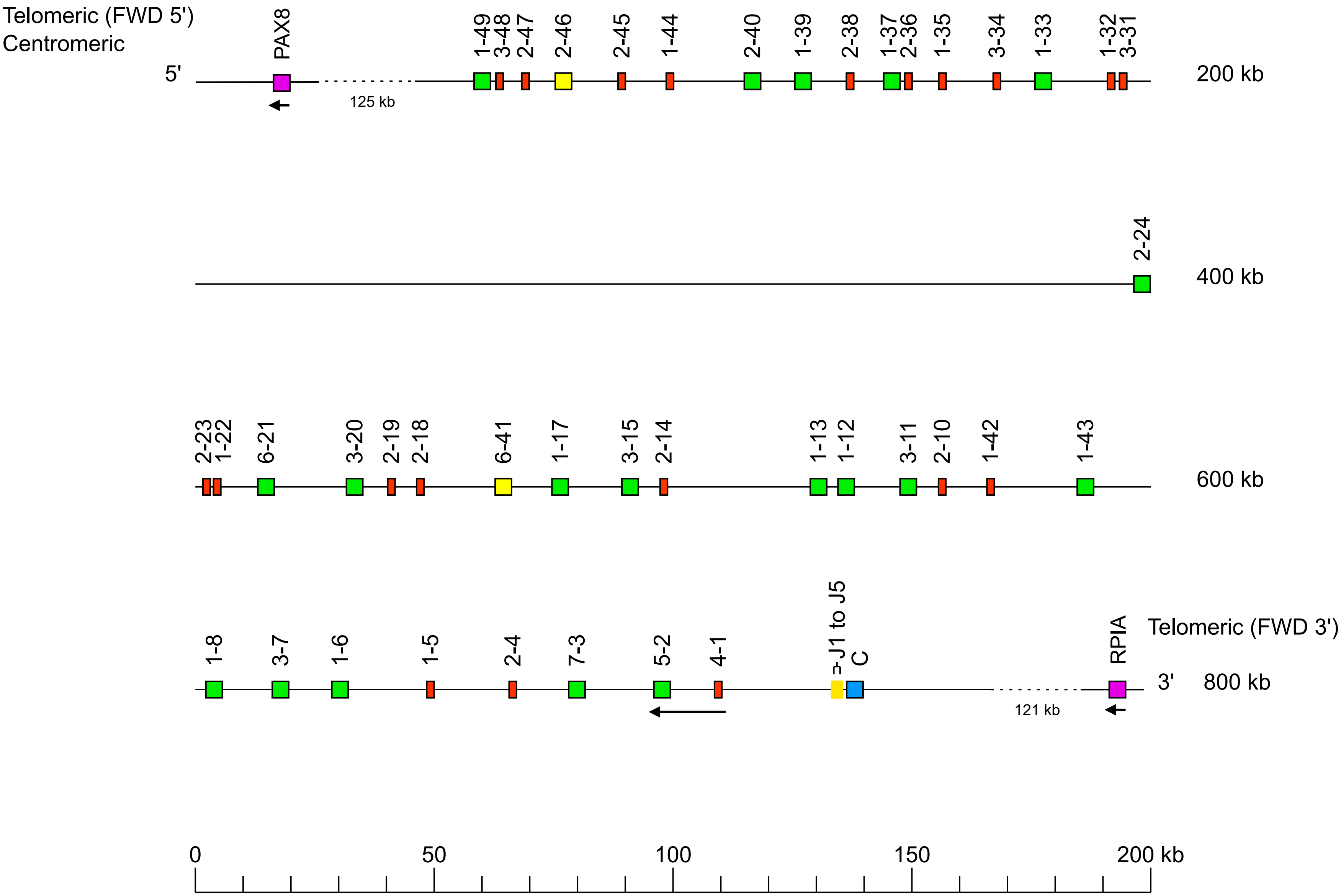

### Supp figure 7: Western lowland gorilla (*Gorilla gorilla gorilla*) IGK locus on chromosome 2A assembly NHGRI_mGorGor1-v1.1-0.2.freeze_mat

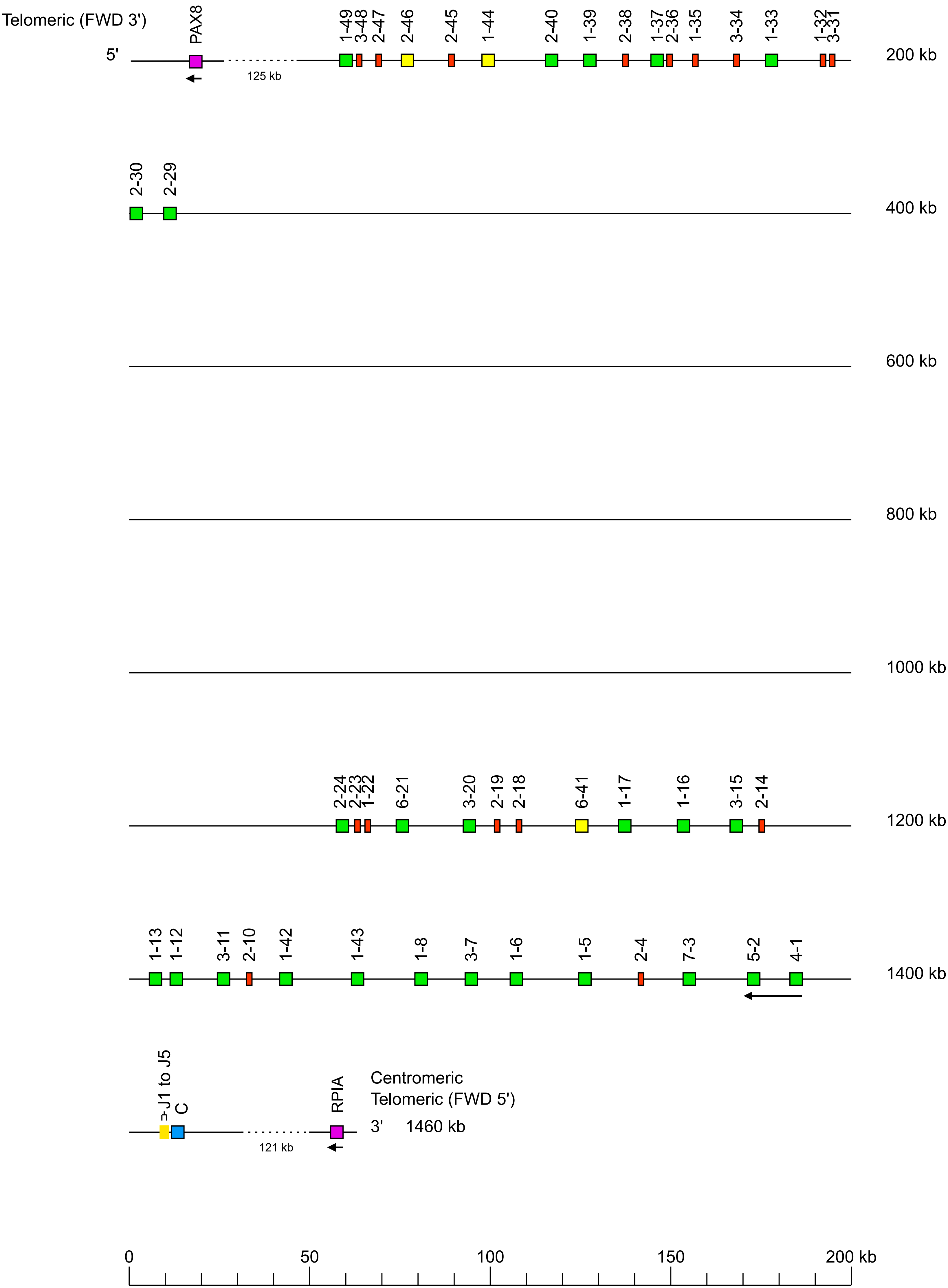

**Supp figure 8: Western lowland gorilla (*Gorilla gorilla gorilla*) IGK locus on chromosome 2A assembly NHGRI_mGorGor1-v1.1-0.2.freeze_pat**

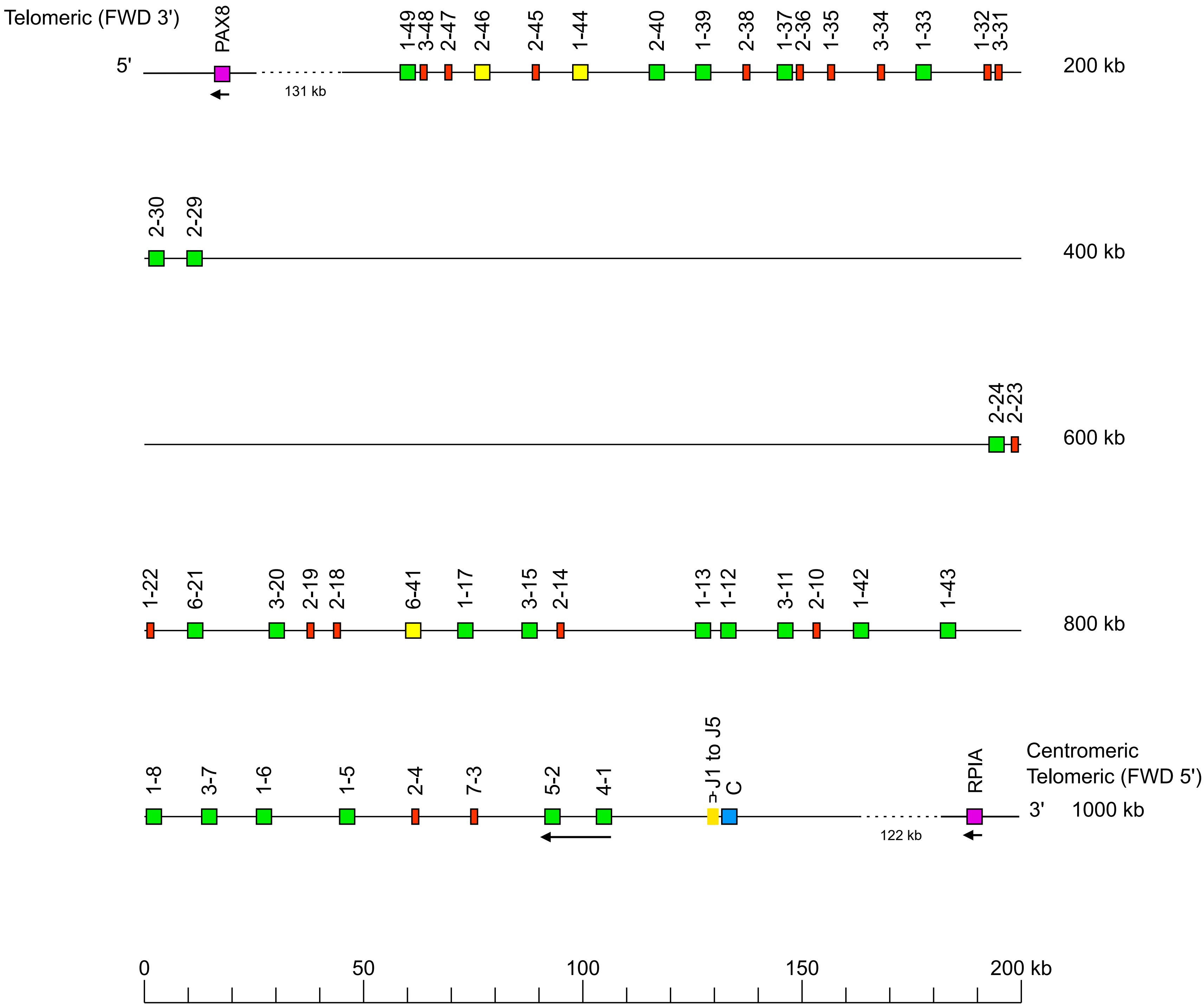
