## Supplemental Figures 9 10 11 12 for "Deciphering of *Gorilla gorilla gorilla* Immunoglobulin Loci in Multiple Genome Assemblies and Enrichment of IMGT Resources"

### Legend

- V-GENE functionnal
- V-GENE ORF
- V-GENE pseudogene
- J-GENE functionnal
- J-GENE ORF
- C-GENE functionnal
- C-GENE pseudogene
- RPI pseudogene
- 5' & 3' IMGT_borne gene functionnal

#### Supp figure 9: Western lowland gorilla (*Gorilla gorilla gorilla*) IGL locus on chromosome 22 assembly Kamilah_GGO_v0

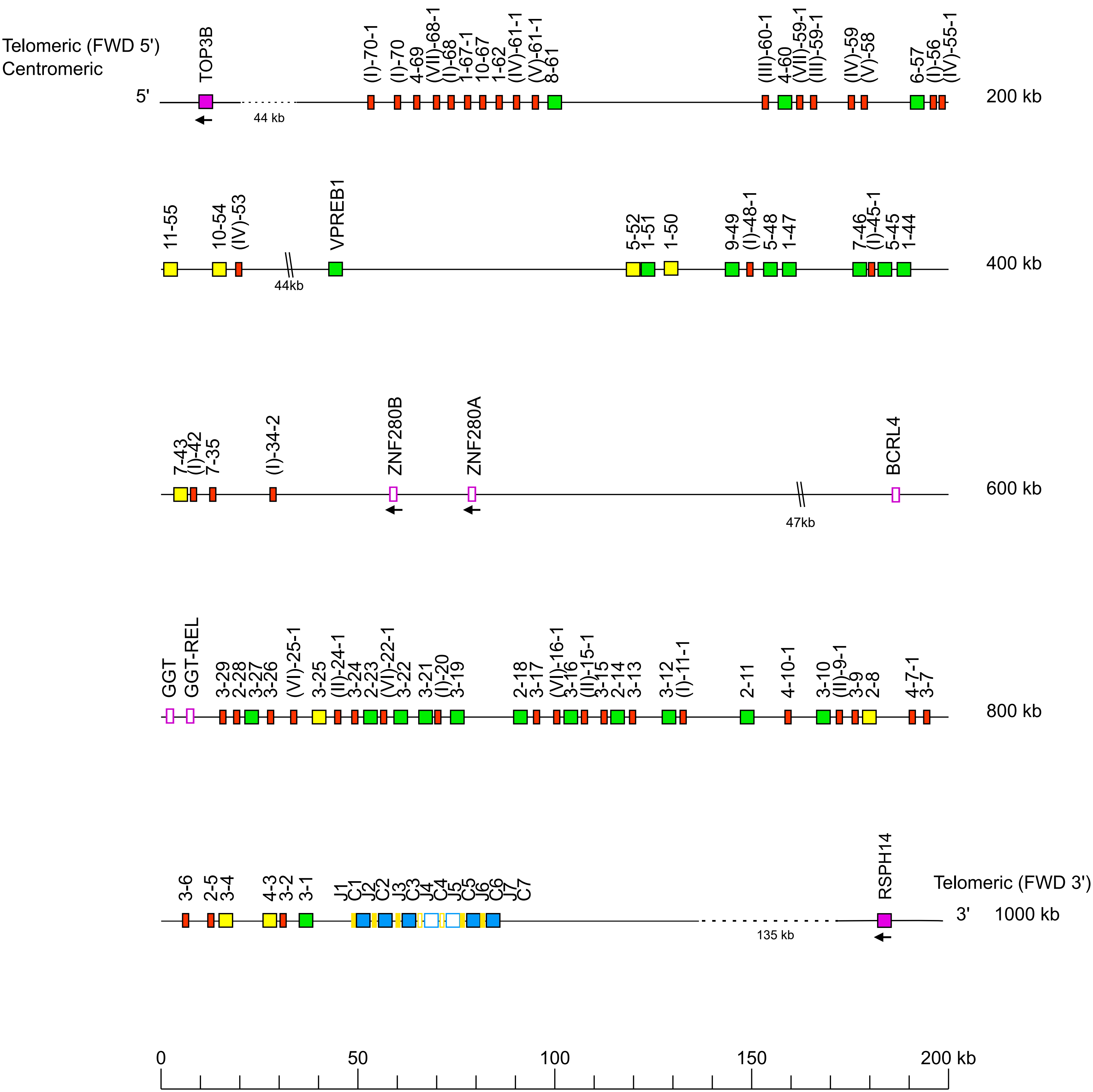

### Supp figure 10: Western lowland gorilla (*Gorilla gorilla gorilla*) IGL locus on chromosome 22 assembly Susie3

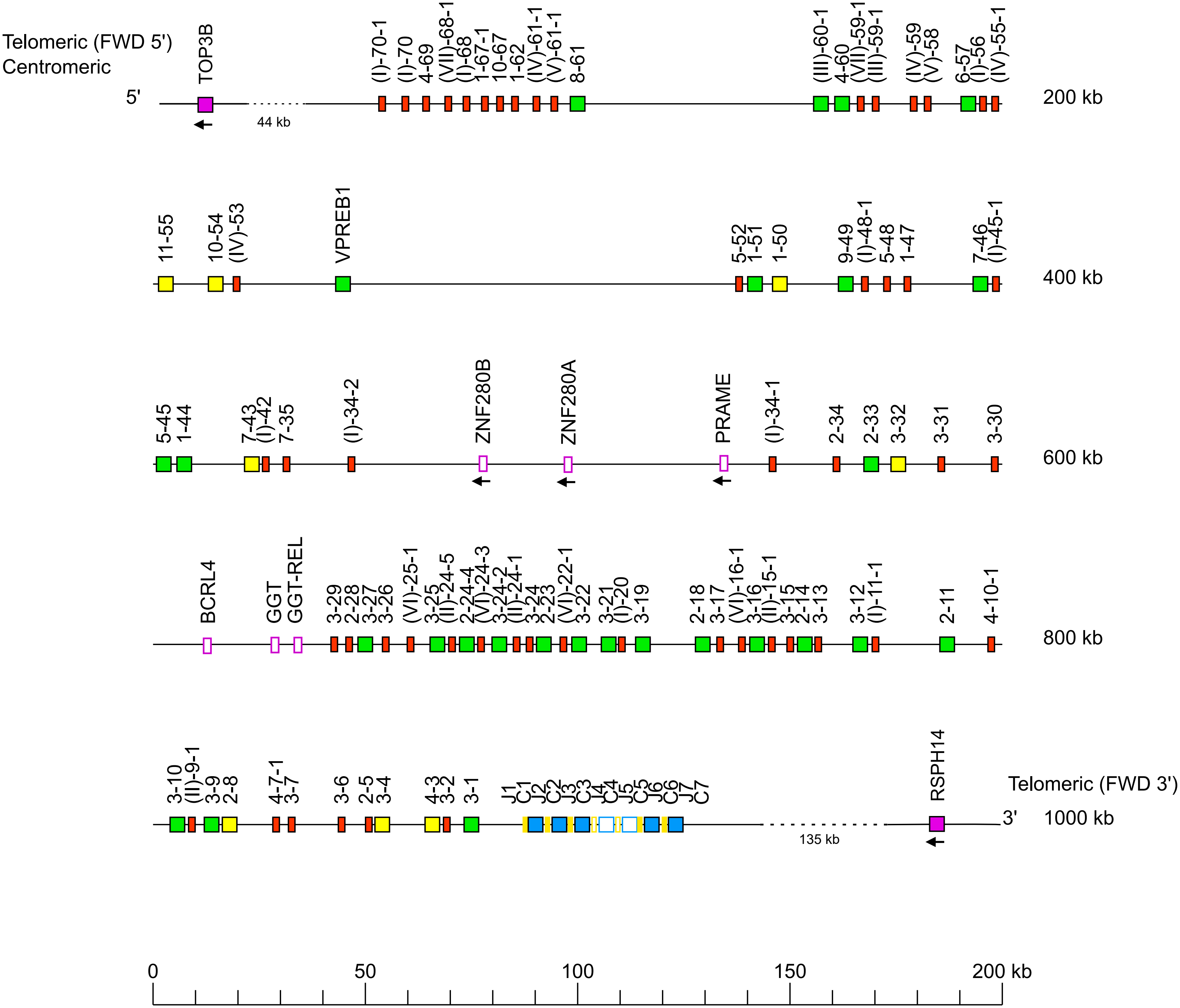

#### Supp figure 11: Western lowland gorilla (*Gorilla gorilla gorilla*) IGL locus on chromosome 22 assembly NHGRI_mGorGor1-v1.1-0.2.freeze_mat

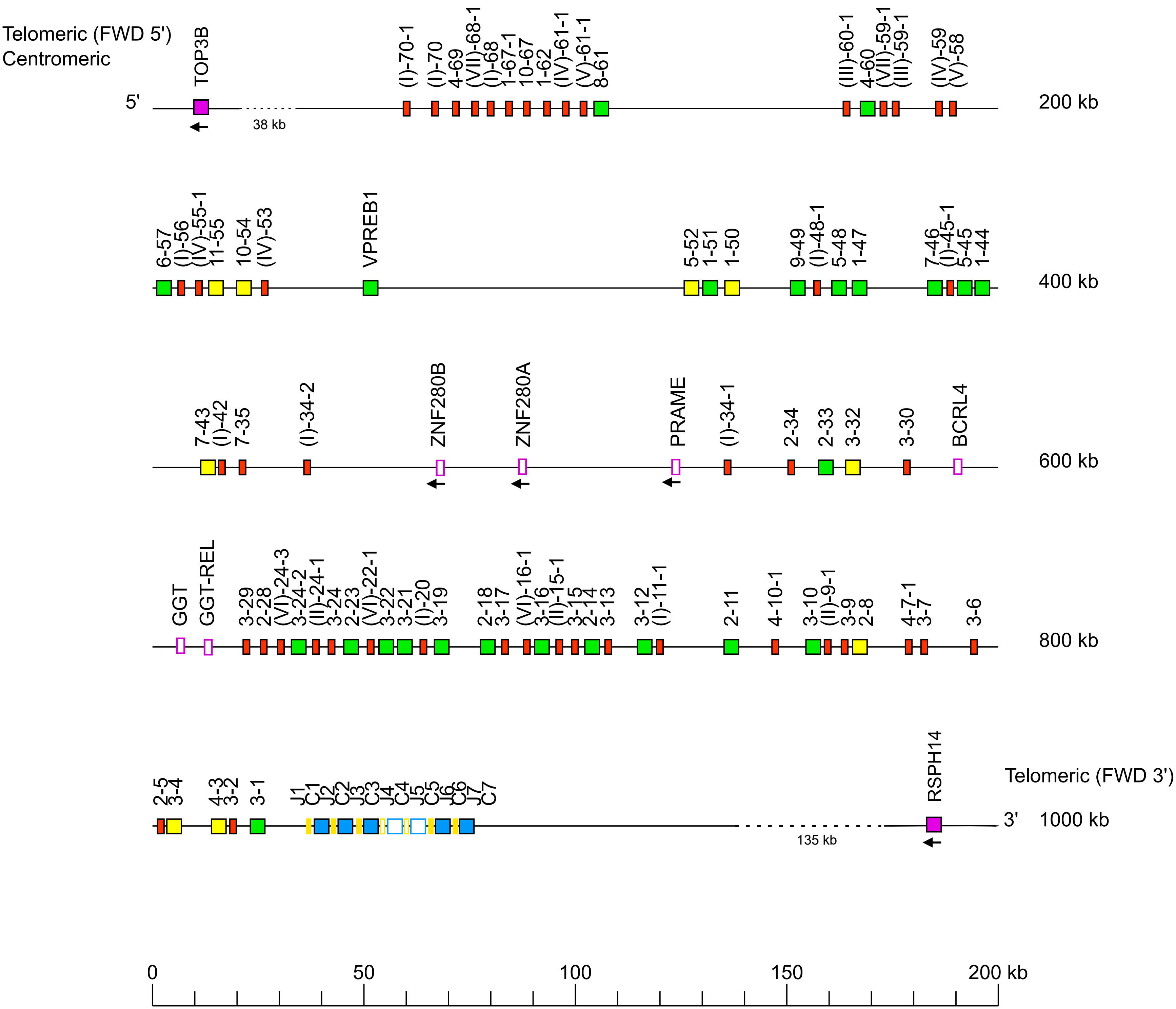

**Supp figure 12: Western lowland gorilla (*Gorilla gorilla gorilla*) IGL locus on chromosome 22 assembly NHGRI_mGorGor1-v1.1-0.2.freeze_pat**

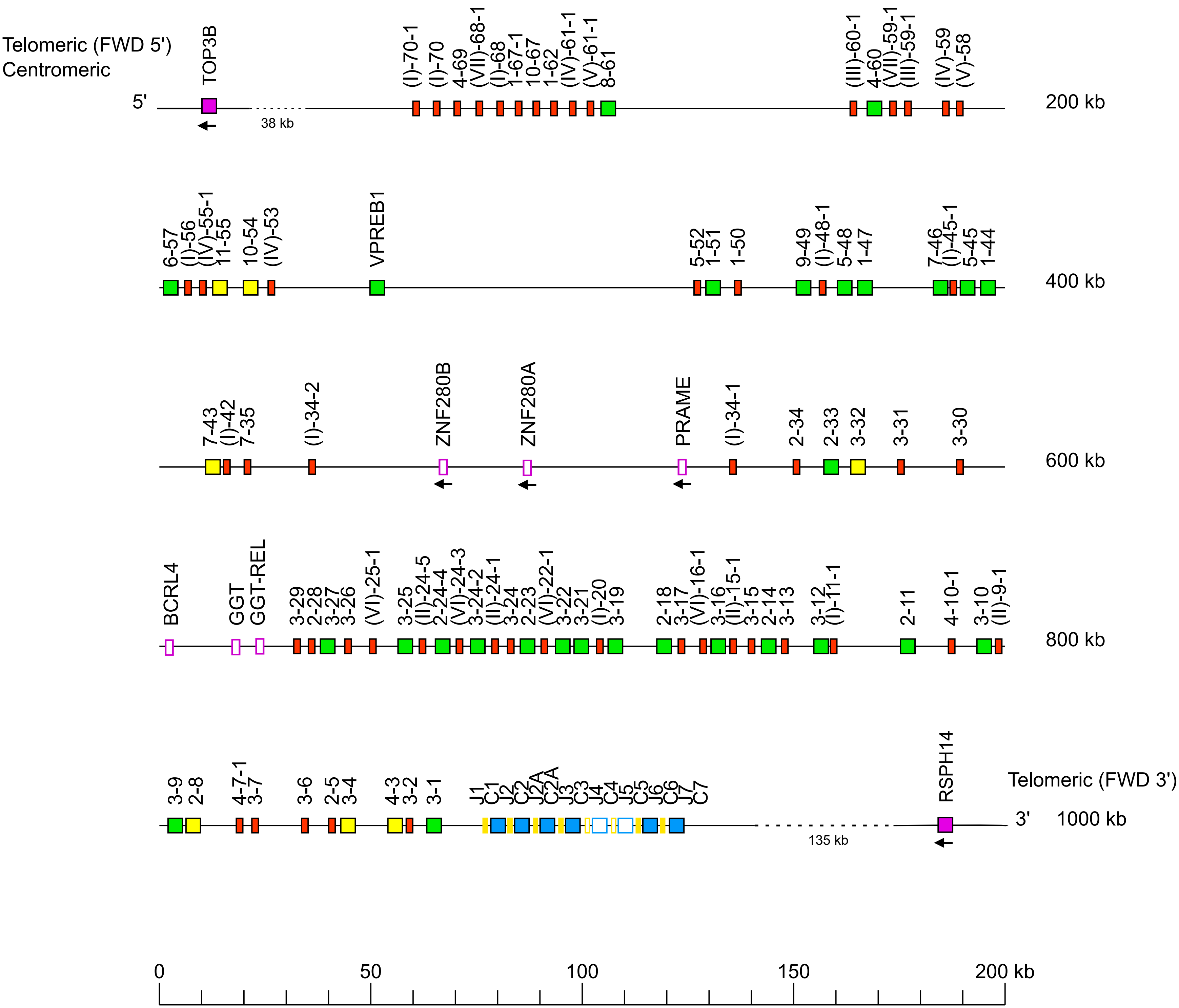
