## Supplemental Figure 13 for "Deciphering of *Gorilla gorilla gorilla* Immunoglobulin Loci in Multiple Genome Assemblies and Enrichment of IMGT Resources"

### Supp figure 13: Phylogenetic tree of all IGK subgroups for *Gorilla gorilla gorilla* and *Homo sapiens*

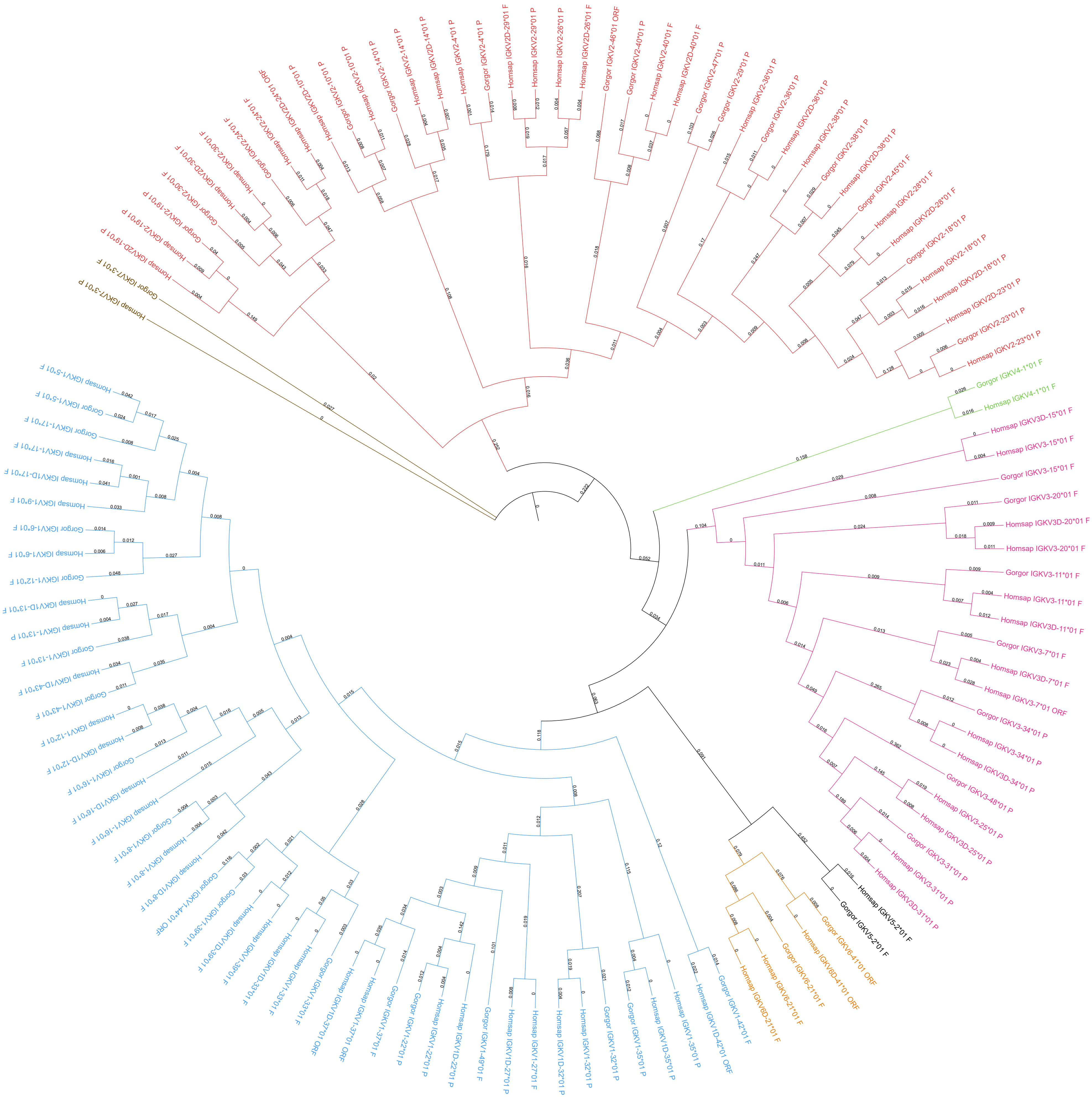
