## Supplemental Figure 14 for "Deciphering of *Gorilla gorilla gorilla* Immunoglobulin Loci in Multiple Genome Assemblies and Enrichment of IMGT Resources"

### Supp figure 14: Phylogenetic tree of all IGL subgroups and clans for *Gorilla gorilla gorilla* and *Homo sapiens*

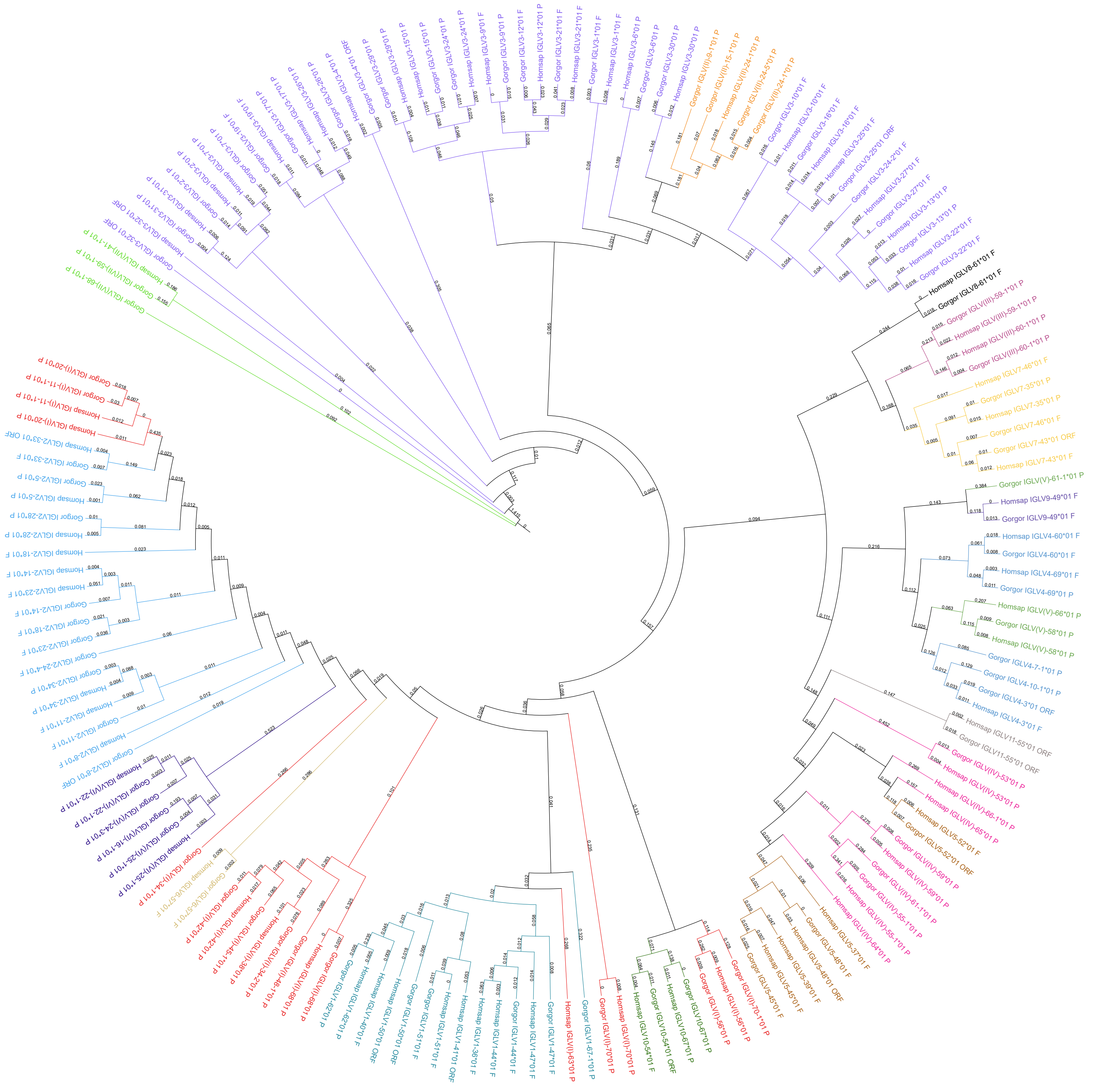
