## Supplemental Figure 15 for "Deciphering of *Gorilla gorilla gorilla* Immunoglobulin Loci in Multiple Genome Assemblies and Enrichment of IMGT Resources"

**Supp figure 15: Number of IGKV genes per IMGT subgroup in human and in the 4 assemblies of *Gorilla gorilla gorilla***

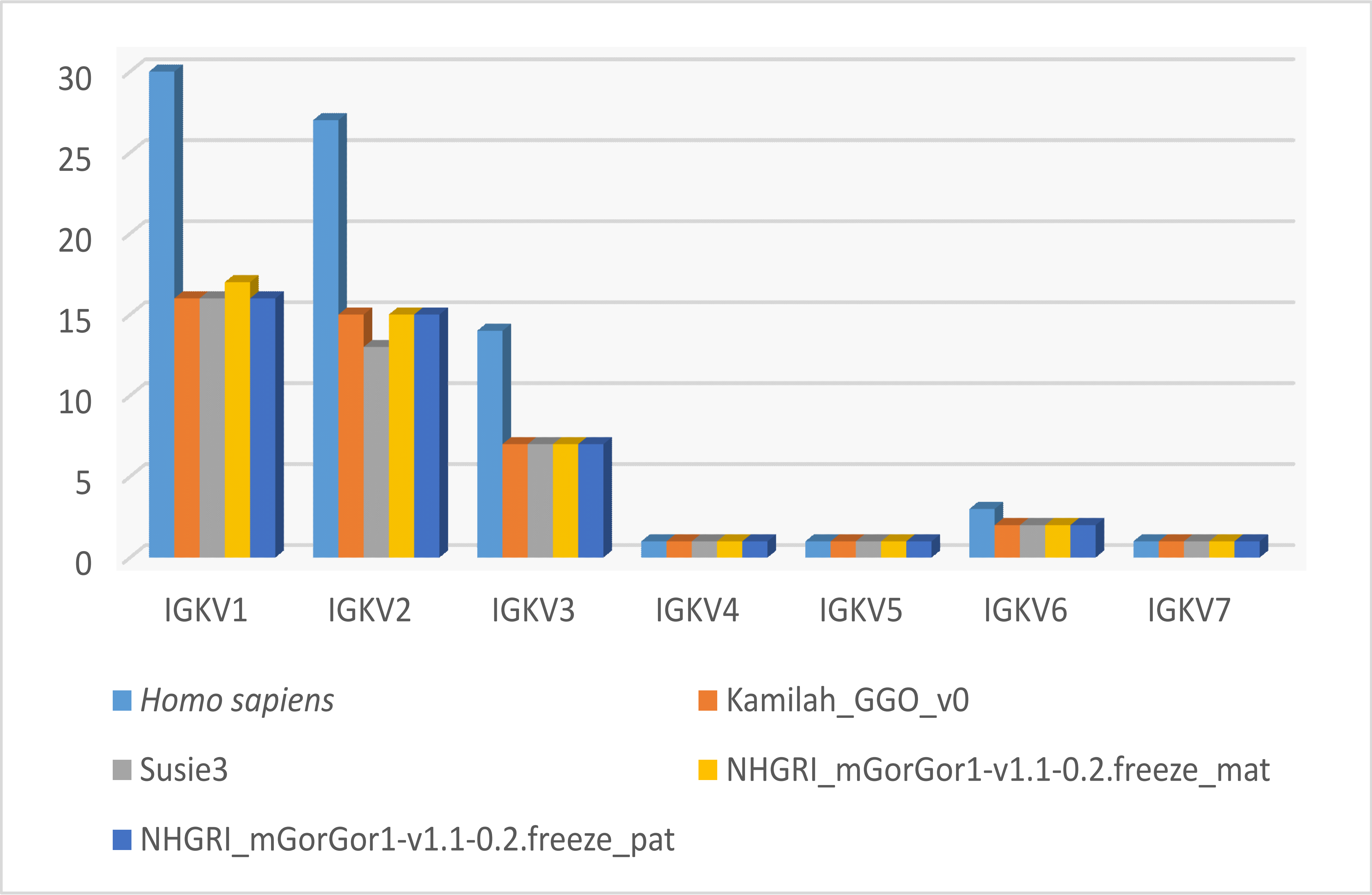
