## Supplemental Figure 16 for "Deciphering of *Gorilla gorilla gorilla* Immunoglobulin Loci in Multiple Genome Assemblies and Enrichment of IMGT Resources"

**Supp figure 16: Number of IGLV genes per IMGT subgroup in human and in the 4 assemblies of *Gorilla gorilla gorilla***

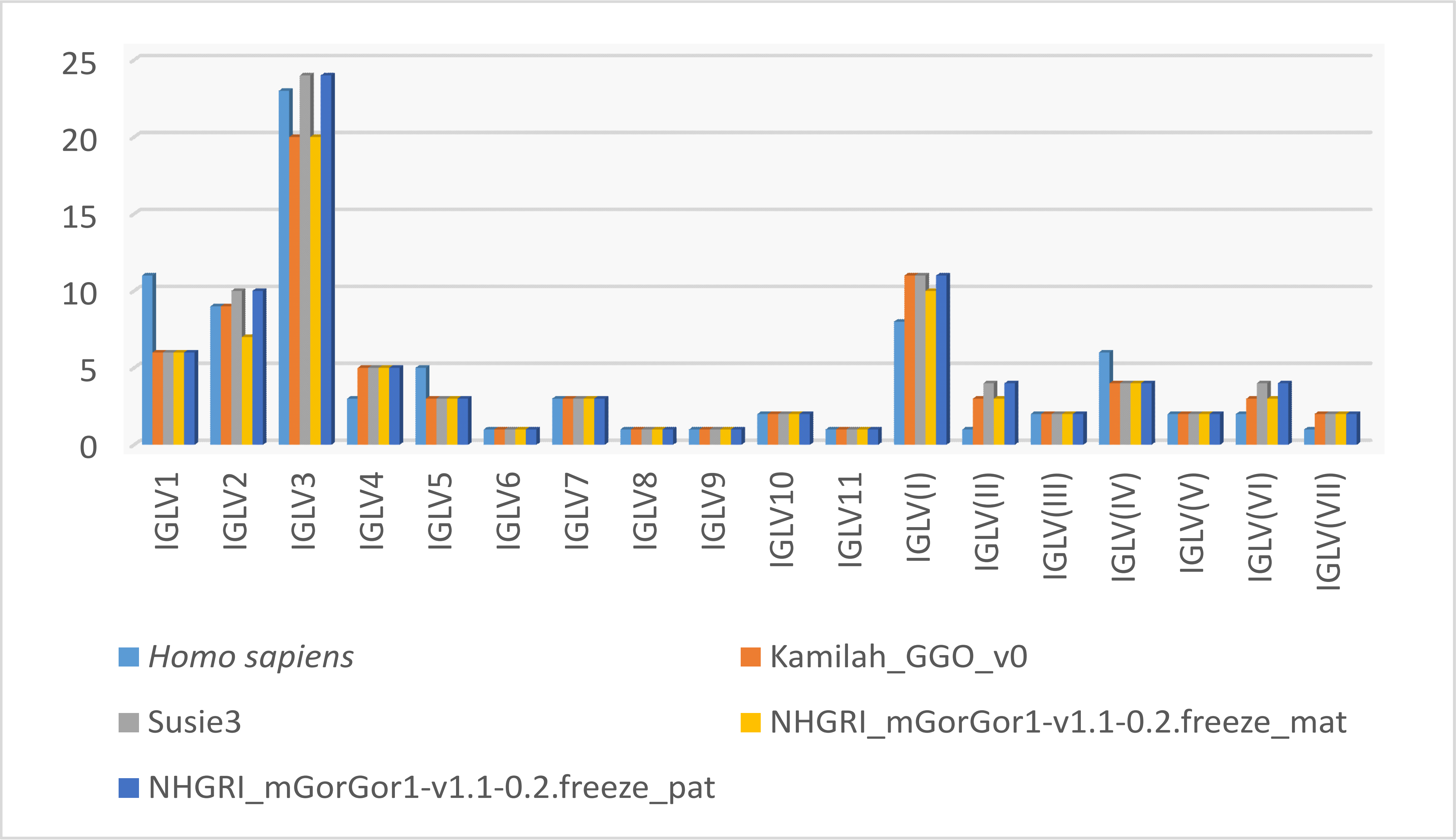
